# Pollinators differentially respond to local and landscape grassland features

**DOI:** 10.1101/2024.07.01.601588

**Authors:** Elinor M. Lichtenberg, Jaclyn Heiser, Kristen A. Baum, John L. Neff, Shalene Jha

**Affiliations:** Department of Biological Sciences and Advanced Environmental Research Institute, University of North Texas, Denton, Texas, 76203-5017, USA; Department of Integrative Biology, The University of Texas at Austin, Austin, TX 78712 USA; AMERMIN, Austin, TX 78759, USA; Department of Integrative Biology, Oklahoma State University, Stillwater, OK 74078, USA; Kansas Biological Survey & Center for Ecological Research, and Department of Ecology and Evolutionary Biology, University of Kansas, Lawrence, KS 66047, USA; Central Texas Melittological Institute, Austin, TX 78731, USA; Lady Bird Johnson Wildflower Center, Austin TX 78739, USA

**Keywords:** biodiversity, habitat, community composition, pollinators, Coleoptera, Diptera, Hymenoptera, Lepidoptera, Great Plains

## Abstract

1. Predicting how habitat composition alters communities of mobile ecosystem service providers remains a major challenge in community ecology. This is partially because separate taxonomic groups that provide the same service may respond uniquely to changes in habitat and associated resource availability. Further, the spatial scale at which habitat features impact each group can vary. Failure to account for these differences significantly limits the ability to quantify shared versus contrasting responses to habitat for important ecosystem service-providing groups.
2. We investigated the impacts of local (habitat patch level) and landscape features in the US Southern Great Plains on groups of pollinating insects with different basic biologies: Coleoptera, Diptera, Hymenoptera, and Lepidoptera. Habitat features included local flower and shelter resources as well as landscape-scale semi-natural habitat.
3. We found that bare ground supported more Hymenoptera and Lepidoptera but fewer Diptera, while more diverse flower communities supported more Hymenoptera but fewer Coleoptera. Interestingly, given that this study occurred in a grassland system, forest cover in the surrounding landscape more strongly affected pollinator diversity than grassland cover did. Landscapes with more woodland had higher Coleoptera and Diptera richness.
4. Our results highlight that pollinator conservation and sustainable land management depend on understanding the habitat needs, including shelter, of diverse pollinators. Because taxa can have opposite responses to specific habitat features or scales, providing a range of grassland management practices (e.g., variety in the timing and type of biomass removal) may be the most effective approach to support the broader pollinator community.

## Introduction

Habitat change is dramatically altering animal biodiversity and its associated ecosystem functions (Raven & Wagner 2021). Despite the importance of mobile animals that provide ecosystem services, such as pollinators and natural enemies (Kremen et al. 2007), past research has often coarsely grouped taxa that provide similar functions. This contrasts with the increasing recognition that species’ basic biologies (e.g., different nesting and foraging behaviors) impact responses to habitat change (Lichtenberg et al. 2017; Rader et al. 2020), even for species that serve similar ecological functions. Indeed, research pioneered with plants highlights that species’ traits can determine both responses to habitat alteration and contributions to ecosystem functions (McGill et al. 2006). Despite this recognition, frameworks for determining community-level response to habitat change often focus on a single taxonomic group. This significantly limits the ability to quantify the shared versus contrasting responses to habitat change for groups that provide important ecosystem services. Advancing this knowledge requires multi-taxa investigation of how distinct groups respond to specific resources and how available those resources are in different habitats.

Insect pollinators are an ideal system for developing a mechanistic understanding of the differential impacts of habitat composition on multi-taxa ecosystem service-providing groups. Insect pollinators are taxonomically diverse, including organisms in the orders Hymenoptera (bees, wasps), Coleoptera (beetles), Diptera (flies), Lepidoptera (butterflies), and others (Rader et al. 2020; Ollerton 2021). Despite this taxonomic breadth, our understanding of how shelter and food resource availabilities affect the abundance and diversity of insect pollinators is often limited to our knowledge of how one taxonomic group (bees, Hymenoptera: Apoidea) responds to floral availability and diversity. Even though butterflies and flies (Lepidoptera & Diptera) can often make up a large fraction of the pollinator community (e.g., Cusser et al. 2016), only a small portion of studies have simultaneously quantified the impacts of habitat variation on these additional orders. Even less research has focused on beetles (Coleoptera), despite their potentially similar role in pollination (Rader et al. 2020; Cusser et al. 2021). These taxonomically diverse insects also provide ecologically and economically valuable pollination services to natural and human-modified habitats (Potts et al. 2016), and many are in decline (Ollerton 2021).

Past work generally finds that pollinator communities respond positively to higher flower availability or diversity (Kral-O’Brien et al. 2021). However, it is not known if this pattern persists across taxonomic groups, especially across environmental conditions. For example, flower availability or diversity promotes bees in prairies and urban gardens (Tonietto et al. 2017; Ballare et al. 2019), butterflies in California mountain meadows (Marschalek et al. 2017), and syrphid flies in German agricultural landscapes (Haenke et al. 2009). However, underlying habitat conditions are dramatically different across these systems. The handful of studies to date that investigated drivers of multiple pollinator taxa within a single system have found positive impacts of flowers on bees, but a mix of positive and no impacts on butterflies, beetles, and flies (e.g., Ekroos et al. 2008, 2013; Fründ et al. 2010; Horak 2014; Gómez-Martínez et al. 2022; He et al. 2022). Indeed, the availability of larval host plants or other feeding substrates can strongly influence these non-bee pollinators (Rader et al. 2020). Given the taxonomic diversity of insect pollinators, and the ecological breadth this diversity encompasses, it is critical to investigate how multiple pollinator groups respond to the same floral resources.

Relative to floral resources, the role of nesting or sheltering habitat in mediating pollinator community composition has not been well studied. Shelter resources such as bare ground, leaf litter, or pithy plant stems are critical for insect survival and reproduction (Kremen et al. 2007), and these shelter needs often vary dramatically among taxa. For example, ground-nesting bees often benefit from bare ground availability (Harmon-Threatt 2020), while beetle and fly species that use leaf litter to hunt for food as larvae or overwinter may benefit from more vegetation cover and less bare ground (e.g., hover flies, Sjödin et al. 2008; pollen beetles, Rusch et al. 2012). In another example, sandy soil often promotes bees but can reduce beetle abundance, while tall vegetation can promote hover flies and beetles but not bees (Sjödin et al. 2008). Determining the responses of different taxa to critical shelter resources is a promising avenue for understanding how entire pollinator communities, not just single taxonomic groups, are expected to respond to habitat change.

Finally, beyond local resources– those found within a habitat patch – such as flowers, nest materials, and shelter sites, the larger surrounding landscape composition can also impact local biodiversity patterns and ecological processes (Tscharntke et al. 2012). These broader landscapes often contain habitat within hundreds of meters of a site and can provide resources that may not be abundant locally. Landscapes can also impact visitation from the regional species pool (Tscharntke et al. 2012) and connect small habitat patches to each other. Several syntheses find that semi-natural habitat in the surrounding landscape can have positive, negative, or no effects on pollinator abundance and diversity (Kennedy et al. 2013; Scheper et al. 2013; Lichtenberg et al. 2017). Responses are hypothesized to be driven partially by mobility differences across taxa due to variation in body size, dispersal behavior, and whether or not a species provisions at a fixed-location nest (central place foraging) (Ballare et al. 2019; Wong et al. 2019; Lichtenberg et al. 2023). Interestingly, larger-scale landscape resources may be less influential when local habitat patches are large or heterogeneous and thereby provide sufficient resources (Tscharntke et al. 2012). Understanding the interactive effects of local- and landscape-scale habitat availability and quality is particularly important when studying an array of taxa, such as multiple orders of insect pollinators.

To understand how distinct taxa within a functional group respond to habitat availability, we investigated the impacts of local and landscape habitat features on insect pollinators that vary in their basic biologies: the orders of bees and wasps (Hymenoptera), butterflies (Lepidoptera), beetles (Coleoptera), and flies (Diptera). These habitat features included multiple local flower and shelter resources and the availability of landscape-level semi-natural habitat. We focused on grasslands in the Southern Great Plains, as this is a highly threatened and understudied habitat (Scholtz & Twidwell 2022) where landscapes are heterogeneous at both small and large scales. Given that the large majority of the insects we studied visit flowers and are thus potential pollinators, for simplicity we refer to these insects as “pollinators”. We addressed two questions. (1) How do pollinators in distinct orders respond to specific local and landscape habitat features? (2) Beyond insect orders, how do local and landscape habitat features structure the larger pollinator community and individual species’ responses? We expected to find differential responses to habitat based primarily on utilized shelter material. For example, we predicted that stem- and cavity-nesting taxa would respond more positively to the amount of woodland in a landscape than ground-nesting taxa. We also expected that central place foragers (most of our Hymenoptera) would be more constrained by local floral resources while other taxa would respond to both local resource availability and landscape-scale habitat heterogeneity.

## Materials and Methods

### Study sites

We conducted our research in June and early July of 2017 at 42 sites located in 11 land units (3-4 sites per land unit) spread across almost 400km of the Cross Timbers ecoregion from central Texas to south central Oklahoma (Fig. 1). Prior to European settlement, this ecoregion historically consisted of grasslands interspersed with dense tree belts, but it has experienced increasing levels of woody encroachment and urbanization (Smith et al. 2019). Further, grasslands in this region are often invaded by two non-native perennial grasses: King Ranch bluestem (*Bothriochloa ischaemum* (Linnaeus) Keng) and Johnsongrass (*Sorghum halepense* (L.) Persoon). These grasses were introduced as cattle forage, but can become very dense and often dominate plant communities after invasion (Fulbright et al. 2013). This may reduce the abundance of flowering forbs and availability of bare ground that many pollinators require for food and nesting (Rader et al. 2020; Kral-O’Brien et al. 2021). This region has an extremely short summer flowering period before conditions become too dry and hot, and we thus could sample each site only once. We ensured that sampling effort on that day matched what was found sufficient in past pollinator studies (e.g., Winfree et al. 2014).

**Figure 1:**
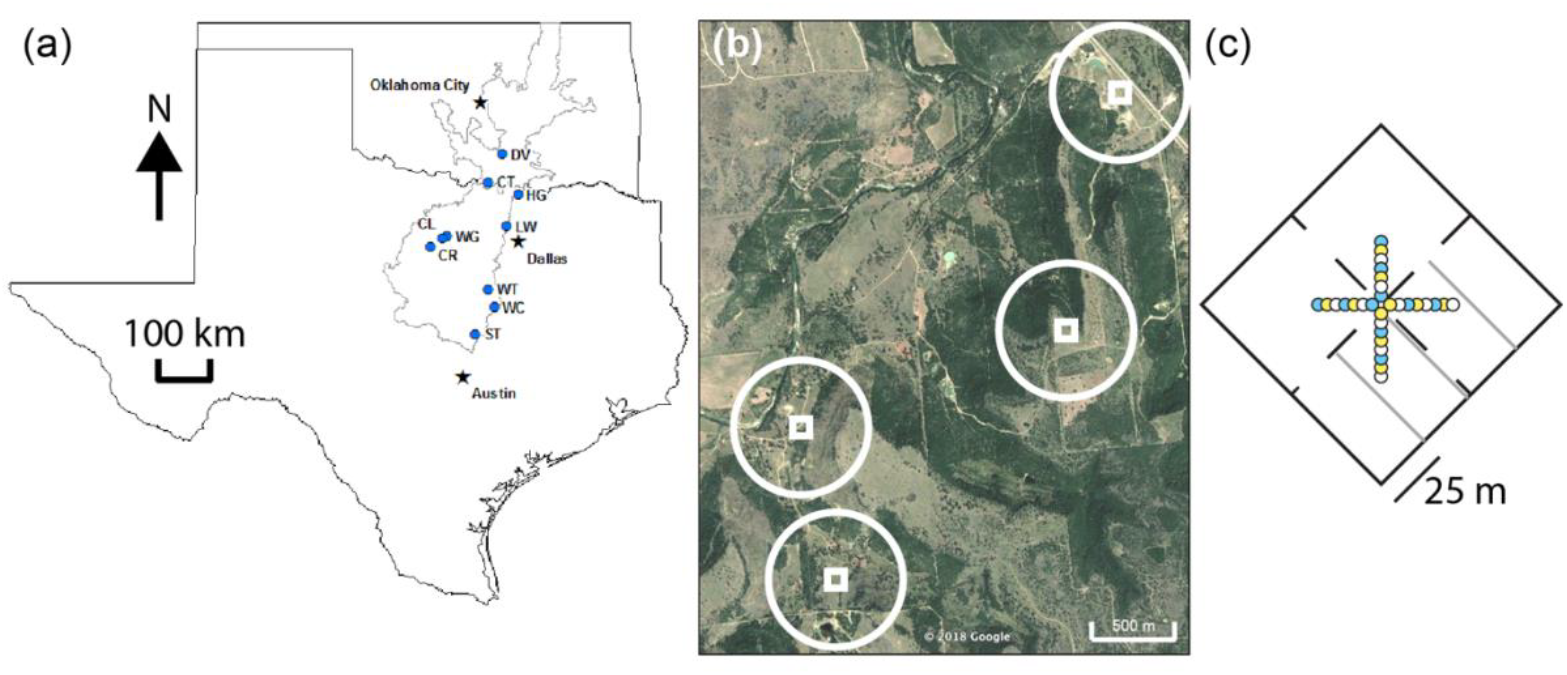
(a) Locations of land units (blue dots) in the Cross Timbers ecoregion (gray outline). (b) Example of a single land unit with four sites (white squares), where circles indicate the landscape within a 1 km radius around each site and squares approximate the plots where we sampled. Plots were centered within sites. (c) To-scale example of a data collection plot, where dashed lines divide quadrants for netting; gray lines are locations of transects for floral, ground cover, and grass sampling; and colored circles are pan traps. Traps were aligned along cardinal directions.

Our sites were a combination of privately owned (4 land units) and managed by the US Army Corps (4 land units), US Fish and Wildlife Service (1 land unit), and Oklahoma Department of Wildlife Conservation (1 land unit). All sites were characterized by grassland plant communities and none had been treated with prescribed fire or seeding in the prior 10 years. Land units were separated by a minimum of 5km (mean: 40.6km, range: 5.1km-69.8km) to the closest unit and sites were separated by a minimum of 0.7km (mean: 1.9km, range: 0.7km-8.3km) within a land unit, which provided spatial independence of the insect communities.

### Insect sampling

We collected field data within a 10,000m^2^ plot (dimensions 100×100m or 50×200m, depending on land configuration) at each site. In each plot, we sampled pollinators through a combination of passive trapping and active netting. We set up 30 pan traps (10 each of bright colors that attract flying insects: white, fluorescent blue, fluorescent yellow; LeBuhn et al. 2003) in a “+” formation at the center of the plot, and separated by 5m for maximum effectiveness (Fig. 1c; Droege et al. 2010). Each trap was a 0.096L soufflé cup (Solo Brand P325) filled with soapy water. We pushed aside vegetation above each trap to ensure they were visible to flying insects. Traps were deployed before 0830 hours Central Daylight Savings Time, and left out for 7 hours (as in Tonietto et al. 2017). On the same day, we collected insects that were visiting flowers in the plot. Four people netted insects observed in the middle of a flower (and thus likely feeding or serving as a pollinator) during morning (between 0900 and 1200 hours) and afternoon (between 1300 and 1600 hours) 30-min sessions (four person hours total per site, including time for insect handling). Netters haphazardly walked through one quarter of the plot (Fig. 1c) and rotated quarters every 15 min to ensure that each netter covered the entire plot across the two netting sessions. Both the trapping and netting methods have previously been successfully implemented in our study region (Ritchie et al. 2016; Ballare et al. 2019).

We stored all trapped specimen in 95% ethanol and netted specimens in the freezer. In the lab, we transferred specimens <2mm long to individual vials of ethanol, while we washed and pinned all other specimens. We then identified each specimen to as fine a taxonomic resolution as feasible, based on morphological characteristics and available keys (e.g., McAlpine 1981; Johnson & Triplehorn 2004; Michener 2007; Schuh et al. 2010; Gibbs et al. 2013; Williams et al. 2014; complete list in Table S1). Specifically, we identified 67% of individuals to species or species group (closely related species that are hard to differentiate), 12% to genus, 4% to family, and 17% to morphospecies. This is similar resolution to past studies covering a large breadth of insect taxa (e.g., Samways et al. 2010; Foster et al. 2019). All specimens are stored in the Jha Lab and will ultimately be deposited in the University of Texas Entomology Collection.

### Floral sampling

We measured flower abundance and richness in 51 1×1m quadrats located along three transects within the plot (following Ritchie et al. 2016; Ballare et al. 2019). These transects ran perpendicular to the 25, 50, and 75m marks along a haphazardly-selected side of the plot for 100×100m plots, and the 50, 100, and 150m marks for the 50×200m plots (Fig. 1c). We placed the quadrat every 3m along these transects, where we identified all plants in bloom and counted the number of flowering inflorescences (e.g., individual flowers, heads, or racemes, depending on species) of each species (as in Ritchie et al. 2016). We then used these data to determine the total inflorescence abundance and richness for a site across all 51 quadrats.

### Ground cover and grass sampling

In the same 51 quadrats, we measured percent cover of the following variables by quantifying the portion of 16 squares (each 25×25cm) in each quadrat, to the nearest half square: bare ground, living grassy or herbaceous vegetation, dead grassy or herbaceous vegetation, rocks, and wood (e.g., tree trunks, large branches on the ground). We used these data to calculate the mean proportion of a 1×1m quadrat that contained each habitat feature (as in Ballare et al. 2019). In two diagonal corners of each quadrat, we also recorded the presence or absence of the invasive perennial grasses: King Ranch bluestem and Johnsongrass. We used these data to calculate the proportion of all observed corners that contained these grasses (invasive grass cover) at a site.

### Quantifying landscape context

We used the 2016 National Land Cover Database to determine the amount of grassland and woodland within a 1km radius of each site (landscape context). First, we quantified in Fragstats (McGarigal et al. 2012) the proportion of this area that consisted of each land cover type. We then summed categories within each habitat to determine the total proportion of the area that consisted of grassland (land cover types: Grassland/Herbaceous and Pasture/Hay) and woodland (land cover types: Deciduous Forest, Evergreen Forest, Mixed Forest, Shrub/Scrub, and Woody Wetlands). Grassland and woodland were the primary land uses around each study site, making up an average 53% and 31% of the overall cover, respectively. These were followed by agriculture (11%), water (8%), developed land (6%), and wetland (2%).

### Data analysis

We investigated how local and landscape habitat availability impacted abundance, diversity, and community composition of insect pollinators in separate orders, conducting all analyses in R (R Core Team 2018). First, we calculated abundance, richness, and evenness (E_var_, Smith & Wilson 1996) of each order at each site (summarized in Table S2). Because there were relatively few Lepidoptera, we analyzed presence rather than abundance, richness, or evenness. We then built negative binomial (abundance data, which were overdispersed) (MASS package, Venables & Ripley 2002), logistic (Lepidoptera presence), Poisson (Diptera richness due to heteroscedasticity), or linear (all other richness and evenness data)(lme4 package, Bates et al. 2015) mixed models with abundance, richness, evenness, or presence of a given order as the response and land unit as a random effect. The random effect controlled for sites within a land unit being geographically close to each other and under similar recent and historic management. To account for heteroscedasticity and an influential outlier (> 4 standard deviations away from the mean), we log-transformed Diptera evenness data and excluded the outlier site from the Diptera abundance regressions, respectively.

For all models, we selected five fixed-effect predictors that captured a range of pollinator habitat needs (e.g., Kremen et al. 2007; Harmon-Threatt 2020) within our sites: floral richness, landscape context, the floral richness-landscape context interaction, bare ground cover, and invasive grass cover (Table S2). Initial analyses confirmed that these variables were not collinear (VIF < 1.25 in all regressions). We did not include floral abundance because it correlated with flower richness (S=5677.97, *r*=-0.57, *p*<0.0001; Table S3) and preliminary analyses confirmed it was less informative. We scaled floral richness for the analyses because all other predictors were proportions. We measured landscape context as the proportion of the surrounding landscape that was grassland or woodland. Because these proportions are highly correlated (S=23186.00, *r*=-0.75, *p*<0.0001; Table S3), we only included one per model, and ran separate models with each landscape type. We then used information-theoretic model selection to determine the best fit model(s) for each metric and insect order (MuMIn, Bartón 2018), selecting the models with low AICc values (within 2 of the lowest value; Burnham & Anderson 2002) and focusing our results on these models. We also calculated rarefied richness and evenness, and estimated richness (Chao1, Gotelli & Colwell 2011). These estimates were highly correlated with measured richness and evenness (Table S4) and thus we focus on measured richness and evenness (as in Fortel et al. 2014).

We then investigated impacts of the same set of predictors on community composition for each order using PERMANOVA (vegan package, Oksanen et al. 2019). To do so, we categorized each predictor variable into “high” (above the median) and “low” (below the median) values. Analyses included site as a grouping variable and removed singleton species to reduce effects of rare taxa. For Coleoptera, we excluded six sites with fewer than two individuals. We assessed significance of each term, including the interaction between flower richness and landscape context, using marginal effects. As with regressions, we ran separate analyses with landscape context as grassland and woodland.

For predictors that significantly affected community composition, we determined species that are strongly associated with a given type of habitat using indicator species analysis (indicspecies package, De Cáceres & Legendre 2009) with phi coefficient correction to account for unequal numbers of sites across groups (Tichý & Chytrý 2006). This provides insight into species’ habitat preferences (De Cáceres & Legendre 2009). To enable indicator species analysis of the interactive effect between flower richness and landscape context, we created an additional variable that had one level for each combination of high and low flower richness and landscape context. Prior to performing indicator species analysis, we ensured homogeneity of multivariate dispersions (vegan package, Oksanen et al. 2019).

## Results

Overall, our sites varied considerably in the local and landscape habitat variables as well as abundance and diversity of insect orders recorded (Table S2). We collected 4499 individuals in 172 unique insect taxa (species, genus, family, or morphospecies) across all sites (Table S5, including taxonomic affiliations and authorities). This included 728 Coleoptera in 25 taxa, 1799 Diptera in 47 taxa, 1891 Hymenoptera in 87 taxa, and 81 Lepidoptera in 13 taxa. Ninety percent of Hymenoptera (1700 individuals) were bees, and the rest were wasps. The most common species were *Ravinia* spp. flies (690 individuals), honey bees (401 individuals), *Epicauta sericans* beetles (342 individuals), *Asyndetus* spp. flies (288 individuals), *Condylostylus* spp. flies (233 individuals), *Lasioglossum coactus* bees (203 individuals), a *Lasioglossum* morphospecies (139 individuals), and *Bombus pensylvanicus* bumble bees (112 individuals). The large majority of individuals and taxa, including those collected only in pan traps, are known to visit flowers (Table S5).

### Abundance and diversity

Regressions indicated that local habitat, particularly bare ground cover and floral richness, strongly impacted pollinator communities (Tables S6-S14). Higher bare ground cover led to significantly higher Hymenoptera abundance (Fig. 2a; Table S12) and Lepidoptera presence (Fig. 2c; Table S15), but lower Diptera abundance (Fig. 2d; Table S9) and Hymenoptera evenness (Fig. 2b; Table S14). Increased floral richness led to greater Hymenoptera abundance and richness (Fig. 3a,b; Tables S12, S13), but lower Coleoptera abundance (Fig. 3c; Table S6). Coleoptera evenness was also higher at sites with more flower species (Fig. 3d). This increase was greater in landscapes with less grassland in one of three best models (Table S8).

**Figure 2:**
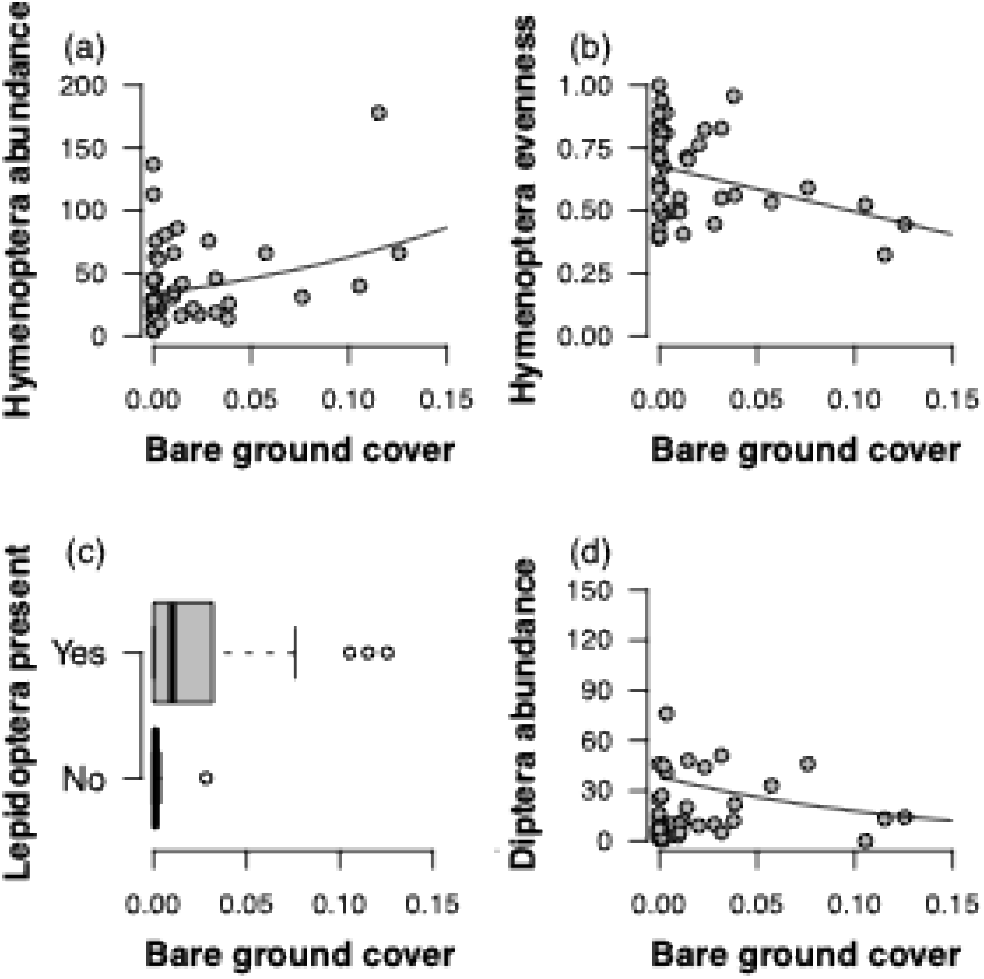
Relationships between bare ground cover (average proportion cover in quadrats) at a site and (a) Hymenoptera abundance, (b) Hymenoptera evenness, (c) Lepidoptera presence/absence, and (d) Diptera abundance. Fitted lines are from negative binomial (a,d) or general linear (b) regressions.

**Figure 3:**
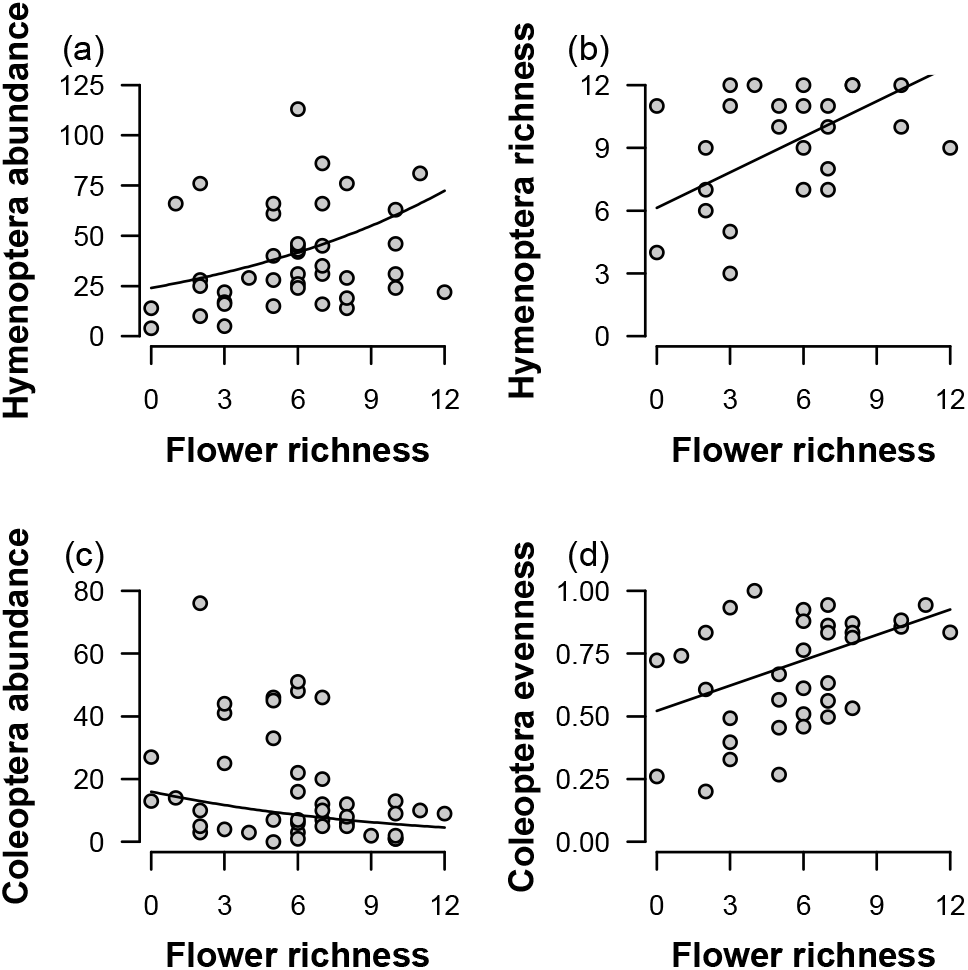
Relationships between flower richness at a site and (a) Hymenoptera abundance, (b) Hymenoptera richness, (c) Coleoptera abundance, and (d) Coleoptera evenness. Fitted lines are from negative binomial (a,c) or general linear (b,d) regressions.

At the landscape scale, woodland cover impacted pollinators more than grassland cover did. Sites in landscapes with more woodland had higher Coleoptera richness (Fig. 4a; Table S7), lower Coleoptera evenness (Fig. 4c; Table S8), and lower Diptera richness (Fig. 4b; Table S10). Sites with greater grassland cover also had greater Coleoptera evenness in one of three best models (Table S8).

**Figure 4:**
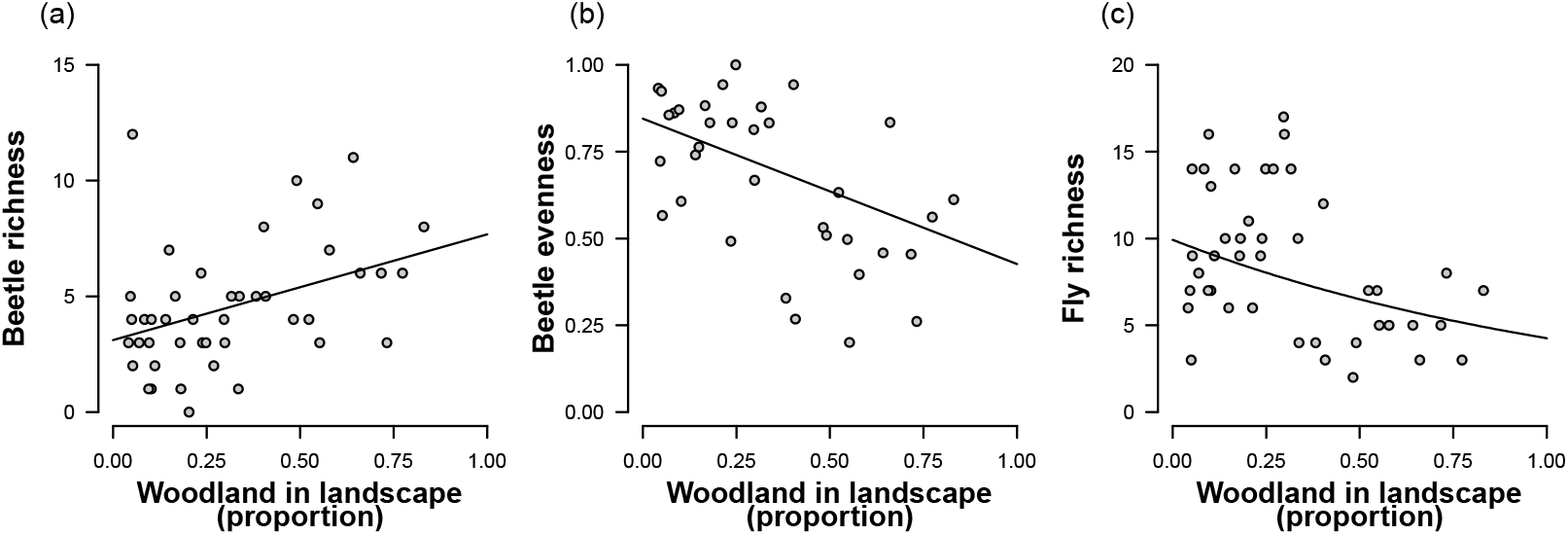
Relationships between the proportion of woodland in the landscape around a site and (a) Coleoptera richness, (b) Coleoptera evenness, and (c) Diptera richness. Fitted lines are from negative binomial (a,c) or Poisson (b) regressions.

Cover of the two invasive grass species, King Ranch bluestem and Johnsongrass, did not correlate with floral richness (Spearman’s test: r= 0.18, p=0.25) and did not impact most pollinator response variables in our regressions. Coleoptera evenness was higher at sites with more invasive grasses in one of six best models (Table S8).

### Community composition

Flower richness and bare ground (local habitat), and proportion of woodland in the landscape, were also important predictors of insect pollinator community composition. Indicator species analyses identified several Diptera and Hymenoptera species that were associated with specific habitat types (Tables S16-S21). *Condylostylus* long-legged flies, *Helicobia rapax* flesh flies, and Sciaridae dark-winged fungus gnats were more likely to be found at sites with little bare ground, while *Geron* bee flies were more likely to occur at sites with more bare ground and, separately, landscapes with more woodland. *Atherigona orientalis* tomato flies were more likely to be found in landscapes with less woodland. Several Diptera and Hymenoptera species, such as *Anoplius nigritus* spider wasps and *Asyndetus* flies, were more likely to be found at specific combinations of local floral richness and landscape woodland availability (Table 1; Fig. S1). We sampled each site only once and thus these patterns could potentially reflect changes across the season. However, most indicator species were found across multiple sites throughout the sampling period (Table S5).

**Table 1:** Habitat associations of indicator taxa. (D) indicates that the taxon is a Diptera, and (H) that it is a Hymenoptera. We indicate nesting location for bees, since these data are available in the literature: (G) indicates ground nesting and (W) indicates wood nesting. Body sizes are mean ± standard deviation inter-tegular distances measured from specimens in our collections. (Nesting references are: (1) Cane et al. 2007; (2) Laport & Minckley 2012; (3) Williams et al. 2014; (4) Danforth et al. 2019; (5) Grab et al. 2019; (6) Kilpatrick 2020; (7) Carril & Wilson 2021)

|  | Low woodland | High woodland |
| --- | --- | --- |
| **Low flower richness** | *Augochlorella aurata* (H; G<sup>5</sup>, 1.72 ± 0.20 mm)<br>*Chrysotus* spp. (D; 0.57 ± 0.03 mm)<br>*Lasioglossum coreopsis* (H; G<sup>7</sup>, 1.04 ± 0.11 mm)<br>*Melissodes communis* (H; G<sup>4</sup>, 3.41 ± 0.11 mm) | *Chrysotus* spp. (D; 0.57 ± 0.03 mm)<br>*Geron* spp. (D; 1.44 ± 0.03 mm)<br>*Melissodes communis* (H; G<sup>4</sup>, 3.41 ± 0.11 mm)<br>*Epimelissodes petulca* (H; G<sup>6</sup>, 3.41 ± 0.13 mm)<br>*Xylocopa virginica* (H; W<sup>2</sup>, 6.34 ± 0.20 mm) |
| **High flower richness** | *Anoplius nigratus* (H; 1.33 ± 0.03 mm)<br>*Asyndetus* spp. (D; 0.45 ± 0.03 mm)<br>*Geron* spp. (D; 1.44 ± 0.03 mm)<br>*Melissodes communis* (H; G<sup>4</sup>, 3.41 ± 0.11 mm)<br>*M. tepaneca* (H; G<sup>4</sup>, 3.23 ± 0.10 mm)<br>*Osmia subfasciata* (H; W<sup>1</sup>, 2.25 ± 0.10 mm)<br>Tachydromiinae spp. (D; 0.59 ± 0.08 mm)<br>*Tomosvaryella* spp. (D; 0.50 ± 0.04 mm)<br>*Xylocopa virginica* (H; W<sup>2</sup>, 6.34 ± 0.20 mm) | *Bombus fraternus* (H; G<sup>3</sup>, 5.22 ± 0.41 mm)<br>*Chrysotus* spp. (D; 0.57 ± 0.03 mm)<br>*Geron* spp. (D; 1.44 ± 0.03 mm)<br>*Xylocopa virginica* (H; W<sup>2</sup>, 6.34 ± 0.20 mm) |

## Discussion

We used a multi-taxa approach to examine the impacts of local and landscape habitat features on insect pollinators and show that a small number of key habitat features – bare ground cover, floral richness, and landscape-scale forest cover – differentially affected pollinator abundance, richness, evenness, and community composition. Specifically, bare ground promoted Hymenoptera and Lepidoptera, but inhibited Diptera. Further, having a more diverse flower community promoted Hymenoptera but disfavored Coleoptera. Interestingly, given that this study occurred in a grassland system, grassland cover in the surrounding landscape minimally affected insect pollinator communities. Instead, sites with greater forest cover had greater Coleoptera richness, but lower Diptera richness and Coleoptera evenness. Community composition analyses were qualitatively similar to regressions, although landscape composition more strongly impacted community composition than abundance or diversity.

Our results highlight a potential key synergy in management of the two most recognized pollinator orders: Hymenoptera and Lepidoptera. Both benefited from having more bare ground, which was typically distributed in small patches (range: 0 to 13 percent cover of quadrats) and not large swathes. Note that we collected relatively few Lepidoptera and recommend further investigation of bare ground impacts, but our results are consistent with some other studies (e.g., Eilers et al. 2013; Marschalek et al. 2017; but see Dickins et al. 2013; Wilson et al. 2015). Benefits of bare ground for Hymenoptera likely reflect the shelter needs of ground-nesting bees. Ninety percent of our Hymenoptera were bees, and the majority of bees in the study region nest in the ground (Ballare et al. 2019). Studies from other regions have also shown that sites with more bare ground often have more ground-nesting bees (reviewed in Harmon-Threatt 2020). Patches of bare ground can also promote wasps and butterflies (Eilers et al. 2013; Szczepko et al. 2020). In many cases, bare patches have less thatch and thus provide easier nest access for ground-nesting bees and wasps (Brokaw et al. 2023) and perhaps easier access to soil nutrients needed by butterflies that exhibit puddling behavior (Ankola et al. 2021). A second hypothesized explanation for positive response to bare ground is that less vegetative cover creates more favorable microclimates through increased sun exposure. Warmer soil (Anderson & Harmon-Threatt 2016) and vegetation (via heat reflected off the ground; e.g., Eilers et al. 2013; Dickins et al. 2013) can enhance larval development of bees and butterflies, respectively. Third, the visual contrast of plants against bare ground may provide easier visual landmarks for bee nest locations (reviewed in Harmon-Threatt 2020) or facilitate prey detection by predators, which are often more abundant in sites with less vegetation diversity (Lucatero et al. 2024). Finally, in grasslands, bare ground may indicate lower grass dominance and thus greater availability of forb species and the flowers that provide nectar to bees and butterflies (Eilers et al. 2013; Marschalek et al. 2017). While bare ground cover did not correlate with flower abundance or richness in our study (Table S3), reduced competition with grasses may support greater forb access to sunlight and soil nutrients (McCain et al. 2010).

For this same bare ground habitat feature, our results highlight the potential for tradeoffs with dipterans. Specifically, we found fewer Diptera at sites with more bare ground. This pattern was not driven by syrphids, the most commonly-studied Diptera pollinators, which comprised only 1% (20 individuals) of our dipterans. Many Diptera adults are free living while larvae are often aquatic or parasitic (McAlpine 1981). Dolichopodid (34% of our specimens) larvae may also live in other humid environments such as thick mulch (Gill et al. 2011; Cicero et al. 2017) or under bark (Kautz & Gardiner 2019). This suggests that sites with more bare ground, and thus sparser vegetation or leaf litter, may not provide sufficient moist microhabitats for fly larvae, thereby limiting populations of flies that visit flowers as adults. In support of this hypothesis, our community composition analyses found that *Condylostylus* species (Dolichopodidae) that require humid environments were found mainly at sites with less bare ground, while *Geron* species that are Lepidoptera parasitoids (Yeates & Greathead 1997) were found mainly at sites with more bare ground. However, natural histories and local or landscape drivers of fly communities remain understudied (Cicero et al. 2017; Kautz & Gardiner 2019), especially compared to bees and butterflies. Our documentation of this key trade-off between the habitat drivers of Diptera relative to Hymenoptera and Lepidoptera highlights the urgent need to better study flies to advance more comprehensive insect conservation strategies. This includes sampling sites multiple times within a season and across multiple years.

Similar to the divergent order-level responses to ground cover, we found contrasting responses to local-scale floral diversity. Hymenoptera exhibited a positive response to floral richness while Coleoptera exhibited a negative response, though they did exhibit higher evenness in the higher floral richness sites. Bees depend heavily on flowers for both larval and adult food; past studies have similarly found that bee abundance and richness increase with floral richness (Venturini et al. 2017; Kral-O’Brien et al. 2021). A diverse pool of available flowers facilitates diet balancing by floral generalists (Leach & Drummond 2018) and also provides the necessary host plants for floral specialists that collect pollen from only a small number of plant species (Isbell et al. 2017). Flower-rich sites also tend to provide floral availability across the entire growing season (Kremen et al. 2007; Kral-O’Brien et al. 2021). Floral richness may be particularly important in grasslands such as our study sites that have not been under intensive management, where there were fewer flowers than in managed prairies (Griffin et al. 2021). Beetle responses to floral diversity are less understood. The several studies that investigate this relationship find positive (Gómez-Martínez et al. 2022) or no (Sjödin et al. 2008; Toivonen et al. 2022) responses. This lack of consensus may reflect the small number of studies focusing on beetle pollinators or beetles’ broad range of larval food resources (Johnson & Triplehorn 2004).

At the landscape scale, we found two notable patterns. First, although we sampled in open grasslands, our insect communities were more strongly affected by the amount of woodland in the landscape than the amount of grassland. Trees provide resources such as cavities for larval development or overwintering, flowers (especially early in the year), and larval food (Rusch et al. 2012; Söderman et al. 2016; Mola et al. 2021). Consistent with this, our community composition analyses found *X. virginica* carpenter bees, which make holes in trees to house their nests, were largely in sites surrounded by more woodland. These bees exhibit signatures of high natal site fidelity within a few hundred meters, despite being capable of long-distance flight (Ballare & Jha 2021). Further, 33% of our beetle specimens, representing 72% of the taxa we found, were in families known to be associated with trees: Anthribidae, Buprestidae, Cerambicidae, Curculionidae, and Mordellidae (Irmler et al. 2010). On the other hand, the observed decrease in fly richness in landscapes with more woodland suggests that many fly species rely largely on open rather than forested habitat for meeting their resource and habitat needs. Understanding the mechanisms behind this pattern, which differs from syrphids’ reliance on forested landscapes (e.g., Sjödin et al. 2008; Söderman et al. 2016), is hindered by a lack of research on the non-syrphid fly species that formed the large majority of our individuals, and on landscape impacts on flies in general (Senapathi et al. 2017; Rader et al. 2020), and requires further research.

Additionally, while some studies find that landscape context modulates local habitat impacts on insect communities (e.g., Batáry et al. 2011; Lichtenberg et al. 2017), in our system the two scales acted relatively independently. Interactive impacts of landscape heterogeneity and local management or habitat availability on insect communities have largely been investigated in cropping systems (Senapathi et al. 2017; but see, e.g., Griffin et al. 2021; Lane et al. 2022). Landscape interactions with local conditions may be less important in areas dominated by semi-natural habitat where resources are overall more abundant and movement among habitat patches is less restricted. Theoretically, landscape moderation of field-scale management in agricultural systems is expected to occur mainly in landscapes that consist of less than 20% non-crop habitat (Tscharntke et al. 2012). In contrast, the landscapes around our sites were very homogeneous, with 50-100% semi-natural habitat. Thus, movement patterns resulting from central place foraging may not be mediated by landscape context in this system. Consistent with movement being constrained at a small spatial scale due to animals’ biologies, we found that Hymenoptera, many of which are constrained by the need to provision a fixed-location nest, responded minimally to landscape composition.

Finally, while we found two invasive grass species – King Ranch bluestem and Johnsongrass – across a number of our sites and often at high densities, their cover did not impact pollinator abundance or richness. These grasses often dominate plant communities after invasion, reducing availability of the forbs that pollinators depend on (Fulbright et al. 2013). While little research has explicitly investigated impacts of invasive grasses on pollinators (Stout & Morales 2009), there is some evidence that these grasses can reduce invertebrate (Fulbright et al. 2013) or bee (Pei et al. 2023) abundance and diversity, and that invasive grass removal can promote butterflies (Gowdy et al. 2022). The present study did not find evidence that invasive grasses were depressing forbs, as invasive grass cover did not correlate with flower abundance or richness at our study sites. However, our study focused on floral metrics and did not examine the effect of invasive grasses on the entire (flowering and non-flowering) plant community, which may be negatively impacted by invasive grasses (e.g., Perkins & Hatfield 2014). Additional research is required to evaluate reproductive versus vegetative impacts of invasive grasses on native plants, especially those native species which are critical for provisioning floral resources for pollinator communities.

Overall, our study highlights the need to understand how multiple habitat features, at the local and landscape scales, impact pollinators across distinct taxonomic groups. We showed that responses to these habitat features vary systematically both among insect orders and across individual species. Further, our results indicate that the composition of semi-natural habitat, at both local and landscape scales, has differential impacts on members of the pollinator community. Managing systems for diverse pollinator communities necessitates meeting the resource needs of insects that have sometimes opposite responses to habitat features such as bare ground or floral richness. Thus, regimes that provide a range of grassland management practices (e.g., variety in the timing and type of biomass removal) may be a more effective approach to support the broader pollinator community than implementing a single management practice at a single point in time. We also highlight that data-driven pollinator conservation requires more research to understand the responses of a broader array of pollinator taxa and to determine which habitat features are most critical to manage.

## Acknowledgements

We thank the US Army Corps of Engineers, Hagerman National Wildlife Refuge, the Oklahoma Department of Wildlife Conservation, and private landowners for providing access to field sites. We thank Llewyn Blossfeld, Shannon Dang, Bridget Harter, Katie Strain, and Sam Wilhelm for field and lab assistance; Llewyn Blossfeld, Parrish Brady, Eliana Buenaventura, Val Bugh, Shannon Dang, Nicholas Fensler, Jason Hansen, Elizabeth Lopez, Julia Mlynarek, James Pitts, Mike Quinn, Brian Rabar, Ed Riley, Tín Rodriguez, Anna Solecki, and Alex Wild for insect identification; Dan Levine for GIS assistance; Michael Caballero for fly measurements; and two anonymous reviewers for feedback. This project was funded by US Fish and Wildlife Service Competitive State Wildlife Grant #492118/494242.

