## Supplementary material for "Pollinators differentially respond to local and landscape grassland features": Table S

**Table S1: Taxonomic resources used to identify insects.**

| **Order** | **Additional taxonomic information** | **Source(s)** |
|---|---|---|
| All | | (Johnson & Triplehorn 2004) |
| Diptera | | (McAlpine 1981) |
| Diptera | Bombyliidae | (Kits et al. 2008) |
| Diptera | Syrphidae | (Miranda et al. 2013) |
| Diptera | Tachinidae | (O'Hara 2013) |
| Diptera | Tephritidae | (White 1988) |
| Coleoptera | *Epicauta* | (Pinto 1991) |
| Hymenoptera | Anthophila (bees) | (Mitchell 1960, 1962; Michener 2007) |
| Hymenoptera | *Agapostemon* | (Roberts 1972) |
| Hymenoptera | *Augochlorella* | (Coelho 2004) |
| Hymenoptera | *Bombus* | (Williams et al. 2014) |
| Hymenoptera | *Calliopsis* | (Shinn 1967) |
| Hymenoptera | *Ceratina* | (Daly 1973) |
| Hymenoptera | Chalcidoidea | (Grissell & Schauff 1990) |
| Hymenoptera | *Heriades* | (Michener 1938) |
| Hymenoptera | *Hylaeus* | (Snelling 1970) |
| Hymenoptera | *Lasioglossum* | (Gibbs et al. 2013) |
| Hymenoptera | *Megachile* (*Litomegachile*) | (Bzdyk 2012) |
| Hymenoptera | Pompilidae | (Dreisbach 1948) |

**Table S2: Habitat characteristics and insect biodiversity at our sites. Lepidoptera were present at 32 (out of 42) sites. Fly abundance excludes an outlier with 192 individuals.**

| Variable | Min value | Median value | Max value | Regression details table |
|---|---|---|---|---|
| Flower richness | 0 | 6 | 12 | |
| Landscape-scale grassland cover | 0.17 | 0.51 | 0.92 | |
| Landscape-scale woodland cover | 0.04 | 0.25 | 0.83 | |
| Bare ground cover | 0.00 | 0.003 | 0.13 | |
| Invasive perennial grass cover | 0.00 | 0.23 | 1.00 | |
| Coleoptera abundance | 0 | 10 | 76 | S6 |
| Coleoptera richness | 0 | 4 | 12 | S7 |
| Coleoptera evenness | 0.20 | 0.72 | 1.00 | S8 |
| Diptera abundance | 4 | 33 | 134 | S9 |
| Diptera richness | 2 | 8 | 16 | S10 |
| Diptera evenness | 0.24 | 0.50 | 0.94 | S11 |
| Hymenoptera abundance | 4 | 31 | 178 | S12 |
| Hymenoptera richness | 3 | 12 | 22 | S13 |
| Hymenoptera evenness | 0.32 | 0.59 | 1.00 | S14 |
| Lepidoptera presence | NA | NA | NA | S15 |

**Table S3:** Correlations among habitat variables, determined via Spearman's rank correlation tests.

| **Measure 1** | **Measure 2** | **Test statistic (S)** | **r** | ***p*-value** |
|---|---|---|---|---|
| Flower abundance | Flower richness | 5677.97 | 0.57 | < 0.0001 |
| Flower abundance | Landscape-scale grassland cover | 14938.13 | -0.13 | 0.4137 |
| Flower abundance | Landscape-scale woodland cover | 15733.19 | -0.19 | 0.2275 |
| Flower richness | Landscape-scale grassland cover | 15038.92 | -0.14 | 0.3862 |
| Flower richness | Landscape-scale woodland cover | 12496.62 | 0.06 | 0.7193 |
| Bare ground cover | Flower abundance | 15116.39 | -0.14 | 0.3658 |
| Bare ground cover | Flower richness | 11153.68 | 0.16 | 0.3121 |
| Bare ground cover | Landscape-scale grassland cover | 16600.85 | -0.25 | 0.1010 |
| Bare ground cover | Invasive grass cover | 14914.86 | -0.13 | 0.4202 |
| Bare ground cover | Landscape-scale woodland cover | 8206.69 | 0.38 | 0.0119 |
| Landscape-scale grassland cover | Landscape-scale woodland cover | 23186.00 | -0.75 | < 0.0001 |
| Invasive grass cover | Flower abundance | 11645.66 | 0.12 | 0.4408 |
| Invasive grass cover | Flower richness | 10862.14 | 0.18 | 0.2485 |
| Invasive grass cover | Landscape-scale grassland cover | 16202.80 | -0.22 | 0.1498 |
| Invasive grass cover | Landscape-scale woodland cover | 12865.81 | 0.03 | 0.8558 |

**Table S4:** Spearman's correlations between abundance and measured, rarefied, and estimated (Chao1) richness and evenness values within each insect order.

| Measure 1 | Measure 2 | Test statistic (S) | r | *p*-value |
|---|---|---|---|---|
| Coleoptera abundance | Coleoptera evenness | 13290.19 | -0.86 | <0.0001 |
| Coleoptera abundance | Coleoptera rarefied evenness | 412.07 | -0.13 | 0.6672 |
| Coleoptera abundance | Coleoptera rarefied richness | 441.11 | -0.21 | 0.4872 |
| Coleoptera abundance | Coleoptera richness | 2850.39 | 0.78 | <0.0001 |
| Coleoptera abundance | Coleoptera estimated richness | 4233.88 | 0.66 | <0.0001 |
| Coleoptera evenness | Coleoptera rarefied evenness | 36.00 | 0.90 | <0.0001 |
| Coleoptera evenness | Coleoptera rarefied richness | 8.00 | 0.98 | <0.0001 |
| Coleoptera evenness | Coleoptera richness | 9369.85 | -0.31 | 0.0678 |
| Coleoptera evenness | Coleoptera estimated richness | 8726.11 | -0.22 | 0.1996 |
| Coleoptera rarefied evenness | Coleoptera rarefied richness | 20.00 | 0.95 | <0.0001 |
| Coleoptera rarefied evenness | Coleoptera richness | 77.63 | 0.79 | 0.0014 |
| Coleoptera rarefied evenness | Coleoptera estimated richness | 186.00 | 0.49 | 0.0929 |
| Coleoptera rarefied richness | Coleoptera richness | 57.46 | 0.84 | 0.0003 |
| Coleoptera rarefied richness | Coleoptera estimated richness | 124.00 | 0.66 | 0.0171 |
| Coleoptera richness | Coleoptera estimated richness | 939.13 | 0.92 | <0.0001 |
| Diptera abundance | Diptera evenness | 22430.13 | -0.82 | <0.0001 |
| Diptera abundance | Diptera rarefied evenness | 4394.96 | 0.02 | 0.9071 |
| Diptera abundance | Diptera rarefied richness | 4058.81 | 0.10 | 0.6100 |
| Diptera abundance | Diptera estimated richness | 5231.39 | 0.60 | <0.0001 |
| Diptera abundance | Diptera richness | 3562.15 | 0.73 | <0.0001 |
| Diptera evenness | Diptera rarefied evenness | 1434.00 | 0.68 | 0.0001 |
| Diptera evenness | Diptera rarefied richness | 1778.00 | 0.60 | 0.0005 |
| Diptera evenness | Diptera estimated richness | 15474.81 | -0.25 | 0.1046 |
| Diptera evenness | Diptera richness | 16140.43 | -0.31 | 0.0473 |
| Diptera rarefied evenness | Diptera rarefied richness | 1002.00 | 0.78 | <0.0001 |
| Diptera rarefied evenness | Diptera estimated richness | 2709.82 | 0.40 | 0.0298 |

| **Measure 1** | **Measure 2** | **Test statistic (S)** | **r** | ***p*-value** |
|---|---|---|---|---|
| Diptera rarefied evenness | Diptera richness | 2315.13 | 0.48 | 0.0066 |
| Diptera rarefied richness | Diptera estimated richness | 1576.80 | 0.65 | 0.0001 |
| Diptera rarefied richness | Diptera richness | 1300.68 | 0.71 | <0.0001 |
| Diptera estimated richness | Diptera richness | 1052.12 | 0.92 | <0.0001 |
| Hymenoptera abundance | Hymenoptera evenness | 24566.27 | -0.85 | <0.0001 |
| Hymenoptera abundance | Hymenoptera rarefied evenness | 8086.64 | -0.35 | 0.0449 |
| Hymenoptera abundance | Hymenoptera rarefied richness | 7759.23 | -0.30 | 0.0936 |
| Hymenoptera abundance | Hymenoptera richness | 4272.91 | 0.68 | <0.0001 |
| Hymenoptera abundance | Hymenoptera estimated richness | 10412.04 | 0.21 | 0.1686 |
| Hymenoptera evenness | Hymenoptera rarefied evenness | 1526.00 | 0.74 | <0.0001 |
| Hymenoptera evenness | Hymenoptera rarefied richness | 1566.00 | 0.74 | <0.0001 |
| Hymenoptera evenness | Hymenoptera richness | 17094.81 | -0.29 | 0.0586 |
| Hymenoptera evenness | Hymenoptera estimated richness | 12981.94 | 0.02 | 0.8998 |
| Hymenoptera rarefied evenness | Hymenoptera rarefied richness | 694.00 | 0.88 | <0.0001 |
| Hymenoptera rarefied evenness | Hymenoptera richness | 3802.51 | 0.36 | 0.0370 |
| Hymenoptera rarefied evenness | Hymenoptera estimated richness | 4209.41 | 0.30 | 0.0938 |
| Hymenoptera rarefied richness | Hymenoptera richness | 2914.47 | 0.51 | 0.0023 |
| Hymenoptera rarefied richness | Hymenoptera estimated richness | 3109.04 | 0.48 | 0.0047 |
| Hymenoptera richness | Hymenoptera estimated richness | 4019.72 | 0.70 | <0.0001 |

**Table S5:** Abundance of each taxon collected. Hymenoptera morphospecies names are unique taxa that have not yet been formally described. The “Visits flowers” column provides references indicating that a given taxon is known to visit flowers. “This study” indicates that we observed the taxon on flowers during netting sessions. Sampling periods are A: 1-15 June, 17-21 June, 22-28 June, 1-13 July. Each sampling period contains roughly the same number of sites and sampling days.

| **Order** | **Family** | **Scientific name/morphospecies ID** | **Visits flowers** | **Abundance** | **Periods present** |
|---|---|---|---|---|---|
| Coleoptera | Anthribidae | *Trigonorhinus limbatus* (Say) | This study, (Valentine 1998) | 20 | A-C |
| Coleoptera | Buprestidae | *Acmaeodera mixta* Horn | This study, (MacRae & Nelson 2003; Evans 2014) | 85 | All |
| Coleoptera | Buprestidae | *Acmaeodera neglecta* Fall | (MacRae & Nelson 2003; Evans 2014) | 8 | B |
| Coleoptera | Buprestidae | *Acmaeodera neoneglecta* Fisher | (MacRae & Nelson 2003; Evans 2014) | 1 | A |
| Coleoptera | Buprestidae | *Acmaeodera obtusa* Horn | This study, (Evans 2014) | 17 | All |
| Coleoptera | Buprestidae | *Acmaeodera ornatoides* Barr | (Evans 2014) | 13 | A, B |
| Coleoptera | Carabidae | *Anisodactylus* spp. | (Papp & Darvas 1998) | 1 | B |
| Coleoptera | Cerambycidae | *Strangalia sexnotata* Haldeman | This study, (Papp 1984; Evans et al. 2003) | 1 | B |
| Coleoptera | Cerambycidae | *Typocerus octonotatus* (Haldeman) | This study, (Papp 1984) | 4 | D |
| Coleoptera | Chrysomelidae | *Chaetocnema* spp. | (Kaiser-Bunbury & Müller 2009; Honěk et al. 2016) | 13 | A, C, D |
| Coleoptera | Chrysomelidae | *Diabrotica undecimpunctata howardi* Barber | This study, (Lavigne 1976; Papp 1984; Clark et al. 2004; Eben 2022) | 1 | C |
| Coleoptera | Chrysomelidae | *Diachus auratus* (Fabricius) | This study | 1 | C |
| Coleoptera | Chrysomelidae | *Disonycha leptolineata* Blatchley | This study, (Arnett et al. 2002) | 1 | C |

| **Order** | **Family** | **Scientific name/morphospecies ID** | **Visits flowers** | **Abundance** | **Periods present** |
|---|---|---|---|---|---|
| Coleoptera | Chrysomelidae | *Glyptina texana* (Crotch) | This study, (Arnett et al. 2002) | 2 | D |
| Coleoptera | Chrysomelidae | *Gratiana pallidula* (Boheman) | | 13 | B-D |
| Coleoptera | Chrysomelidae | *Helocassis clavata* (Fabricius) | | 1 | D |
| Coleoptera | Chrysomelidae | *Oulema* spp. | (Papp 1984; Clark et al. 2004) | 3 | D |
| Coleoptera | Coccinellidae | *Harmonia axyridis* (Pallas) | This study, (Hagen 1962; Lichtenberg et al. 2023) | 1 | A |
| Coleoptera | Coccinellidae | *Hippodamia convergens* Guerin | This study, (Hagen 1962; Lichtenberg et al. 2023) | 2 | A |
| Coleoptera | Coccinellidae | *Scymnus* spp. | (Hagen 1962) | 1 | A |
| Coleoptera | Curculionidae | Ceutorhynchinae spp. | (Hatt et al. 2018) | 1 | D |
| Coleoptera | Curculionidae | *Cylindrocopturus* spp. | (Arnett et al. 2002) | 2 | C, D |
| Coleoptera | Curculionidae | *Geraeus* sp. 1 | (Arnett et al. 2002) | 1 | D |
| Coleoptera | Curculionidae | *Geraeus* sp. 2 | This study, (Arnett et al. 2002) | 1 | D |
| Coleoptera | Curculionidae | *Odontocorynus* spp. | This study, (Arnett et al. 2002) | 67 | All |
| Coleoptera | Curculionidae | *Rhyssomatus* sp. 1 | This study, (Arnett et al. 2002) | 2 | D |
| Coleoptera | Curculionidae | *Smicronyx* sp. 1 | (Arnett et al. 2002) | 1 | A |
| Coleoptera | Dermestidae | *Cryptorhopalum* group *triste* LeConte | This study, (Kiselyova 2002) | 29 | All |
| Coleoptera | Meloidae | *Epicauta atrata* (Fabricius) | This study, (Pinto 1991) | 5 | B, D |
| Coleoptera | Meloidae | *Epicauta callosa* LeConte | This study, (Pinto 1991) | 29 | All |
| Coleoptera | Meloidae | *Epicauta immaculata* (Say) | This study, (Pinto 1991) | 10 | B, C |
| Coleoptera | Meloidae | *Epicauta sericans* LeConte | This study, (Pinto 1991) | 342 | All |

| Order | Family | Scientific name/morphospecies ID | Visits flowers | Abundance | Periods present |
|---|---|---|---|---|---|
| Coleoptera | Meloidae | *Nemognatha explanata* Enns | This study, (Enns 1955) | 10 | A, C |
| Coleoptera | Meloidae | *Nemognatha lurida* LeConte | This study, (Dillon 1952; Enns 1955) | 1 | C |
| Coleoptera | Meloidae | *Nemognatha piazata* (Fabricius) | This study, (Enns 1955; Werner et al. 1966) | 1 | C |
| Coleoptera | Meloidae | *Pyrota invita* Horn | This study, (Dillon 1952) | 2 | A |
| Coleoptera | Meloidae | *Zonitis perforata* Casey | This study, (Dillon 1952; Enns 1955) | 1 | C |
| Coleoptera | Meloidae | *Zonitis vittigera* (LeConte) | This study, (Dillon 1952; Enns 1955) | 3 | D |
| Coleoptera | Mordellidae | *Mordella* sp. 1 | This study, (Ford & Jackman 1996) | 16 | All |
| Coleoptera | Mordellidae | *Mordella* sp. 2 | This study, (Ford & Jackman 1996) | 3 | B-D |
| Coleoptera | Mordellidae | sp. 3 | (Ford & Jackman 1996) | 1 | A |
| Coleoptera | Scarabaeidae | *Boreocanthon ebenus* (Say) | (Arnett & Thomas 2000) | 9 | A, D |
| Coleoptera | Scarabaeidae | *Euphoria kernii* Haldeman | This study, (Arnett & Thomas 2000; Orozco 2012) | 1 | A |
| Coleoptera | Staphylinidae | *Baeocera* spp. | (Arnett & Thomas 2000) | 1 | D |
| Diptera | Asilidae | *Mallophora orcina* (Wiedemann) | This study, (Larson et al. 2001) | 6 | D |
| Diptera | Asilidae | *Orthogonis stygia* (Bromley) | This study, (Larson et al. 2001) | 1 | C |
| Diptera | Asilidae | *Ospriocerus* spp. | (Larson et al. 2001) | 7 | A, B |
| Diptera | Asilidae | *Psilocurus* spp. | (Larson et al. 2001) | 3 | D |
| Diptera | Asteiidae | *Leiomyza* spp. | This study | 3 | D |

| Order | Family | Scientific name/morphospecies ID | Visits flowers | Abundance | Periods present |
|---|---|---|---|---|---|
| Diptera | Bombyliidae | *Aphoebanthus* sp. 1 | (Lavigne 1976; Yeates & Greathead 1997; Larson et al. 2001) | 1 | B |
| Diptera | Bombyliidae | *Chrysanthrax cypris* (Meigan) | This study, (Yeates & Greathead 1997; Larson et al. 2001) | 11 | A, C, D |
| Diptera | Bombyliidae | *Chrysanthrax edititius* (Say) | This study, (Yeates & Greathead 1997; Larson et al. 2001) | 2 | D |
| Diptera | Bombyliidae | *Exoprosopa meigenii* (Wiedemann) | This study, (Yeates & Greathead 1997; Larson et al. 2001) | 1 | C |
| Diptera | Bombyliidae | *Geron* spp. | This study, (Yeates & Greathead 1997; Larson et al. 2001) | 86 | All |
| Diptera | Bombyliidae | *Hemipenthes sinuosa* Wiedemann | This study, (Yeates & Greathead 1997; Larson et al. 2001) | 7 | A, C, D |
| Diptera | Bombyliidae | *Lepidanthrax coquilletti* Evenhuis & Greathead | (Yeates & Greathead 1997; Larson et al. 2001) | 1 | B |
| Diptera | Bombyliidae | *Lepidophora lepidocera* (Wiedemann) | This study, (Yeates & Greathead 1997; Larson et al. 2001) | 1 | C |
| Diptera | Bombyliidae | *Neacreotrichus diversa* (Coquillett) | This study, (Yeates & Greathead 1997; Larson et al. 2001) | 1 | A |
| Diptera | Bombyliidae | *Poecilanthrax lucifer* (Fabricius) | This study, (Larson et al. 2001) | 2 | B, C |

| Order | Family | Scientific name/morphospecies ID | Visits flowers | Abundance | Periods present |
|---|---|---|---|---|---|
| Diptera | Bombyliidae | *Poecilognathus punctipennis* (Walker) | This study, (Deyrup 1988; Yeates & Greathead 1997; Larson et al. 2001) | 3 | A, B |
| Diptera | Bombyliidae | *Poecilognathus* sp. 1 | This study, (Yeates & Greathead 1997; Larson et al. 2001) | 11 | A, C, D |
| Diptera | Bombyliidae | *Rhynchanthrax* spp. | This study, (Yeates & Greathead 1997; Larson et al. 2001) | 2 | B, C |
| Diptera | Bombyliidae | *Systoechus solitus* (Walker) | This study, (Yeates & Greathead 1997; Larson et al. 2001; Evans et al. 2003) | 3 | A |
| Diptera | Bombyliidae | *Toxophora amphitea* Walker | This study, (Yeates & Greathead 1997; Larson et al. 2001) | 1 | A |
| Diptera | Bombyliidae | *Villa lateralis* (Say) | This study, (Yeates & Greathead 1997; Pascarella et al. 2001; Larson et al. 2001) | 4 | A, C |
| Diptera | Calliphoridae | *Lucilia* spp. | (McAlpine 1981; Pascarella et al. 2001; Larson et al. 2001) | 2 | B, C |
| Diptera | Chironomidae | *Micropsectra* sp.1 | This study, (McAlpine 1981) | 1 | C |
| Diptera | Chloropidae | *Apallates* spp. | (Larson et al. 2001) | 7 | A, C, D |
| Diptera | Chloropidae | *Chlorops* spp. | (Larson et al. 2001) | 23 | A-C |
| Diptera | Chloropidae | *Conioscinella* spp. | (Larson et al. 2001) | 32 | All |

| **Order** | **Family** | **Scientific name/morphospecies ID** | **Visits flowers** | **Abundance** | **Periods present** |
|---|---|---|---|---|---|
| Diptera | Chloropidae | *Hippelates* spp. | (Lavigne 1976; Larson et al. 2001) | 2 | D |
| Diptera | Chloropidae | *Incertella* spp. | (Larson et al. 2001) | 23 | All |
| Diptera | Chloropidae | *Oscinella* spp. | (Larson et al. 2001) | 1 | A |
| Diptera | Conopidae | *Zodion* spp. | This study, (McAlpine 1981; Larson et al. 2001) | 1 | A |
| Diptera | Culicidae | *Culiseta incidens* (Thomson) | This study, (Larson et al. 2001; Johnson & Triplehorn 2004) | 1 | D |
| Diptera | Dolichopodidae | *Asyndetus* spp. | (Robinson & Deyrup 1997) | 290 | A, C, D |
| Diptera | Dolichopodidae | *Chrysotus* spp. | (van Duzee 1924) | 83 | All |
| Diptera | Dolichopodidae | *Condylostylus* spp. | This study, (Pascarella et al. 2001) | 236 | All |
| Diptera | Dolichopodidae | *Pelastoneurus* sp.1 | (Nadia et al. 2013) | 1 | C |
| Diptera | Fannidae | *Fannia pusio* (Wiedemann) | (Taroda & Gibbs 1982) | 2 | D |
| Diptera | Hybotidae | Tachydromiinae spp. | | 68 | D |
| Diptera | Muscidae | *Atherigona orientalis* Schiner | (Evans et al. 2003) | 13 | C, D |
| Diptera | Muscidae | *Caricea erythrocera* Robineau-Desvoidy | This study | 2 | A, D |
| Diptera | Muscidae | *Coenosia (Limosia)* spp. | (Swales 1979) | 1 | D |
| Diptera | Muscidae | *Haematobia irritans* Linnaeus | | 1 | B |
| Diptera | Muscidae | *Musca* spp. | This study | 1 | B |
| Diptera | Muscidae | *Polietes* sp. 1 | | 3 | A, D |
| Diptera | Muscidae | spp. | | 2 | A, C |
| Diptera | Phoridae | Group 1 | (Larson et al. 2001) | 9 | A, D |
| Diptera | Phoridae | Group 2 | (Larson et al. 2001) | 36 | All |

| Order | Family | Scientific name/morphospecies ID | Visits flowers | Abundance | Periods present |
|---|---|---|---|---|---|
| Diptera | Pipunculidae | *Tomosvaryella* spp. | This study, (Larson et al. 2001) | 4 | A, C |
| Diptera | Sarcophagidae | *Arachnidomyia* sp.1 | (McAlpine 1981) | 1 | D |
| Diptera | Sarcophagidae | *Argoravinia modesta* (Wiedemann) | (McAlpine 1981) | 6 | A, C, D |
| Diptera | Sarcophagidae | *Bercaea cruentata* Meigen | (McAlpine 1981) | 2 | A, D |
| Diptera | Sarcophagidae | *Comasarcophaga* spp. | (McAlpine 1981) | 7 | A, D |
| Diptera | Sarcophagidae | *Helicobia rapax* (Walker) | (McAlpine 1981) | 20 | A, C, D |
| Diptera | Sarcophagidae | Miltogramminae spp. | This study, (McAlpine 1981) | 4 | A, C, D |
| Diptera | Sarcophagidae | *Oxysarcodexia* spp. | (McAlpine 1981; Evans et al. 2003) | 1 | B |
| Diptera | Sarcophagidae | *Rafaelia rufiventris* Townsend | (McAlpine 1981) | 1 | D |
| Diptera | Sarcophagidae | *Ravinia* spp. | This study, (McAlpine 1981) | 690 | All |
| Diptera | Sarcophagidae | *Taxigramma heteroneura* (Meigen) | (McAlpine 1981) | 1 | D |
| Diptera | Sarcophagidae | *Titanogrypa* sp.1 | (McAlpine 1981) | 1 | A |
| Diptera | Sarcophagidae | *Tylomyia* spp. | (McAlpine 1981) | 5 | C, D |
| Diptera | Sarcophagidae | *Udamopyga* spp. | (McAlpine 1981) | 1 | D |
| Diptera | Sarcophagidae | spp. | (McAlpine 1981) | 1 | A |
| Diptera | Sciaridae | *Eugnoriste brevirostris* Coquillett | This study | 2 | A |
| Diptera | Sciaridae | spp. | This study | 10 | A, D |
| Diptera | Sepsidae | *Saltella sphondylii* (Schrank) | (Lavigne 1976; Tepedino et al. 2012) | 1 | D |
| Diptera | Stratiomyidae | *Nemotelus kansensis* Adams | This study, (Larson et al. 2001) | 5 | A, B |
| Diptera | Syrphidae | *Copestylum* spp. | This study, (Larson et al. 2001) | 6 | A, B |

| **Order** | **Family** | **Scientific name/morphospecies ID** | **Visits flowers** | **Abundance** | **Periods present** |
|---|---|---|---|---|---|
| Diptera | Syrphidae | *Dioprosopa clavata* (Fabricius) | (Larson et al. 2001) | 1 | C |
| Diptera | Syrphidae | *Eristalis stipator* Osten Sacken | This study, (Larson et al. 2001) | 1 | D |
| Diptera | Syrphidae | *Palpada furcata* (Wiedemann) | This study, (Larson et al. 2001) | 1 | C |
| Diptera | Syrphidae | *Palpada vinetorum* (Fabricius) | This study, (Pascarella et al. 2001; Larson et al. 2001; Evans et al. 2003) | 2 | C |
| Diptera | Syrphidae | *Paragus haemorrhous* Meigen | This study, (Larson et al. 2001) | 1 | D |
| Diptera | Syrphidae | *Toxomerus marginatus* (Say) | This study, (Pascarella et al. 2001; Larson et al. 2001) | 7 | A, D |
| Diptera | Syrphidae | *Trichopsomyia* sp.1 | This study, (Larson et al. 2001) | 1 | D |
| Diptera | Tachinidae | *Archytas* spp. | This study, (McAlpine 1981; Pascarella et al. 2001; Evans et al. 2003) | 1 | C |
| Diptera | Tachinidae | *Cylindromyia intermedia* (Meigen) | This study, (McAlpine 1981) | 5 | All |
| Diptera | Tachinidae | *Distichona* spp. | This study, (McAlpine 1981) | 1 | D |
| Diptera | Tachinidae | Group 1 | This study, (McAlpine 1981) | 1 | A |
| Diptera | Tachinidae | Group 2 | This study, (McAlpine 1981) | 1 | A |
| Diptera | Tachinidae | Group 3 | (McAlpine 1981) | 1 | D |
| Diptera | Tachinidae | *Gymnoclytia* sp. 1 | This study, (McAlpine 1981) | 1 | A |

| Order | Family | Scientific name/morphospecies ID | Visits flowers | Abundance | Periods present |
|---|---|---|---|---|---|
| Diptera | Tachinidae | *Jurinia smithi/pompalis* (Wulp)/(Reinhard) | This study, (McAlpine 1981; Larson et al. 2001) | 1 | D |
| Diptera | Tachinidae | *Phasia* spp. | This study, (McAlpine 1981; Larson et al. 2001) | 4 | A |
| Diptera | Tephritidae | *Zonosemata electa* (Say) | This study, (McAlpine 1981; Larson et al. 2001) | 1 | D |
| Diptera | | sp. | | 1 | A |
| Hymenoptera | Andrenidae | *Andrena rudbeckiae* Robertson | This study, (Michener 2007) | 5 | C |
| Hymenoptera | Andrenidae | *Calliopsis andreniformis* Smith | (Michener 2007) | 5 | A, C |
| Hymenoptera | Andrenidae | *Perdita cambarella* Cockerell | This study, (Michener 2007) | 54 | D |
| Hymenoptera | Andrenidae | *Perdita ignota* Cockerell | This study, (Michener 2007) | 2 | A, D |
| Hymenoptera | Andrenidae | *Perdita xanthisma* Cockerell | (Michener 2007) | 1 | C |
| Hymenoptera | Andrenidae | *Protandrena rugosa* (Robertson) | This study, (Michener 2007) | 1 | A |
| Hymenoptera | Apidae | *Apis mellifera* Linnaeus | This study, (Michener 2007) | 401 | A-C |
| Hymenoptera | Apidae | *Bombus auricomus* (Robertson) | This study, (Michener 2007) | 1 | C |
| Hymenoptera | Apidae | *Bombus fraternus* (Smith) | This study, (Michener 2007) | 4 | A, C |
| Hymenoptera | Apidae | *Bombus griseocollis* (Degeer) | This study, (Michener 2007) | 43 | C, D |
| Hymenoptera | Apidae | *Bombus pensylvanicus* (Degeer) | This study, (Michener 2007) | 112 | A-C |
| Hymenoptera | Apidae | *Centris atripes* Cresson | This study, (Michener 2007) | 2 | A |

| Order | Family | Scientific name/morphospecies ID | Visits flowers | Abundance | Periods present |
|---|---|---|---|---|---|
| Hymenoptera | Apidae | *Ceratina cockerelli* Smith | This study, (Michener 2007) | 1 | A |
| Hymenoptera | Apidae | *Ceratina shinnersi* Timberlake | This study, (Michener 2007) | 10 | C |
| Hymenoptera | Apidae | *Ceratina strenua* Smith | This study, (Michener 2007) | 35 | D |
| Hymenoptera | Apidae | *Diadasia enavata* (Cresson) | This study, (Michener 2007) | 2 | C |
| Hymenoptera | Apidae | *Diadasia rinconis* (Cresson) | This study, (Michener 2007) | 4 | B |
| Hymenoptera | Apidae | *Diadasia* sp. | (Michener 2007) | 1 | C |
| Hymenoptera | Apidae | *Epimelissodes atripes* (Cresson) | This study, (Michener 2007) | 1 | D |
| Hymenoptera | Apidae | *Epimellisodes grandissima* (Cockerell) | This study, (Michener 2007) | 1 | D |
| Hymenoptera | Apidae | *Epimelissodes obliqua* (Say) | This study, (Michener 2007) | 3 | B |
| Hymenoptera | Apidae | *Epimelissodes petulca* (Cresson) | This study, (Michener 2007) | 5 | B, C |
| Hymenoptera | Apidae | *Melissodes communis* Cresson | This study, (Michener 2007) | 18 | All |
| Hymenoptera | Apidae | *Melissodes coreopsis* Robertson | This study, (Michener 2007) | 5 | A-C |
| Hymenoptera | Apidae | *Melissodes tepaneca* Cresson | This study, (Michener 2007) | 28 | All |
| Hymenoptera | Apidae | *Melissodes wheeleri* Cockerell | This study, (Michener 2007) | 3 | A |
| Hymenoptera | Apidae | *Triepeolus simplex* Robertson | This study, (Michener 2007) | 1 | A |

| Order | Family | Scientific name/morphospecies ID | Visits flowers | Abundance | Periods present |
|---|---|---|---|---|---|
| Hymenoptera | Apidae | *Xylocopa virginica* (Linnaeus) | This study, (Michener 2007) | 89 | B, C |
| Hymenoptera | Bethylidae | sp. H | | 1 | D |
| Hymenoptera | Braconidae | Euphorinae sp. F | (Jervis et al. 1993) | 1 | C |
| Hymenoptera | Braconidae | *Glyptapanteles* sp. | (Jervis et al. 1993) | 1 | D |
| Hymenoptera | Braconidae | sp. 1 | (Pascarella et al. 2001) | 1 | A |
| Hymenoptera | Braconidae | sp. 2 | (Pascarella et al. 2001) | 3 | A, C |
| Hymenoptera | Colletidae | *Colletes birkmanni* Swenk | This study, (Michener 2007) | 1 | A |
| Hymenoptera | Colletidae | *Colletes mandibularis* Smith | This study, (Michener 2007) | 7 | A |
| Hymenoptera | Colletidae | *Hylaeus affinis* (Smith) | (Michener 2007) | 1 | D |
| Hymenoptera | Crabronidae | *Astata clypeata* Parker | This study | 2 | A |
| Hymenoptera | Crabronidae | *Astata mexicana* Cresson | This study | 1 | A |
| Hymenoptera | Crabronidae | *Bembix* sp. | This study, (Evans 1970) | 1 | B |
| Hymenoptera | Crabronidae | *Liris argentatus* (Beauvois) | (Pulawski 2024) | 4 | A |
| Hymenoptera | Crabronidae | *Soleriella* sp. 1 | (Williams 1950) | 24 | All |
| Hymenoptera | Crabronidae | *Soleriella* sp. 2 | (Williams 1950) | 2 | B |
| Hymenoptera | Crabronidae | *Sphecius speciosus* (Drury) | This study | 1 | A |
| Hymenoptera | Crabronidae | *Stictia* sp. 1 | This study, (Pulawski 2024) | 1 | B |
| Hymenoptera | Crabronidae | *Tachysphex* sp. 1 | (Evans 1970) | 3 | A, B, D |
| Hymenoptera | Crabronidae | *Tachysphex* sp. 2 | (Evans 1970) | 16 | All |
| Hymenoptera | Crabronidae | *Tachysphex* sp. 3 | (Evans 1970) | 2 | B, D |
| Hymenoptera | Crabronidae | *Tachysphex* sp. 4 | This study, (Evans 1970) | 1 | B |
| Hymenoptera | Crabronidae | *Tachysphex* sp. 5 | (Evans 1970) | 1 | A |

| Order | Family | Scientific name/morphospecies ID | Visits flowers | Abundance | Periods present |
|---|---|---|---|---|---|
| Hymenoptera | Crabronidae | *Tachysphex* sp. 6 | (Evans 1970) | 6 | B-D |
| Hymenoptera | Crabronidae | *Tachytes amazonus* Smith | (Pulawski 2024) | 2 | A, B |
| Hymenoptera | Crabronidae | *Trypoxylon* sp. 1 | (Pulawski 2024) | 4 | A, C, D |
| Hymenoptera | Encyrtidae | sp. j | (Jervis et al. 1993) | 1 | C |
| Hymenoptera | Encyrtidae | sp. | (Jervis et al. 1993) | 1 | C |
| Hymenoptera | Eulophidae | sp. 1 | (Jervis et al. 1993) | 1 | D |
| Hymenoptera | Eulophidae | sp. 2 | (Jervis et al. 1993) | 1 | D |
| Hymenoptera | Eulophidae | sp. 3 | (Jervis et al. 1993) | 1 | B |
| Hymenoptera | Eupelmidae | Eupelminae sp. c | (Jervis et al. 1993) | 1 | A |
| Hymenoptera | Halictidae | *Acunomia nortoni* (Cresson) | This study, (Michener 2007) | 1 | C |
| Hymenoptera | Halictidae | *Agapostemon splendens* (Lepeletier) | This study, (Michener 2007) | 2 | A, D |
| Hymenoptera | Halictidae | *Agapostemon texanus* Cresson | This study, (Michener 2007) | 37 | All |
| Hymenoptera | Halictidae | *Augochlorella aurata* (Smith) | This study, (Michener 2007) | 37 | C, D |
| Hymenoptera | Halictidae | *Augochloropsis metallica* (Fabricius) | This study, (Michener 2007) | 6 | A, D |
| Hymenoptera | Halictidae | *Augochloropsis sumptuosa* (Smith) | This study, (Michener 2007) | 2 | C, D |
| Hymenoptera | Halictidae | *Halictus ligatus* Say | This study, (Michener 2007) | 12 | All |
| Hymenoptera | Halictidae | *Lasioglossum (Dialictus)* sp. | (Michener 2007) | 1 | B |
| Hymenoptera | Halictidae | *Lasioglossum bardum* (Cresson) | (Michener 2007) | 4 | B |
| Hymenoptera | Halictidae | *Lasioglossum bruneri* (Crawford) | This study, (Michener 2007) | 33 | A, B |

| Order | Family | Scientific name/morphospecies ID | Visits flowers | Abundance | Periods present |
|---|---|---|---|---|---|
| Hymenoptera | Halictidae | *Lasioglossum callidum* (Sandhouse) | This study, (Michener 2007) | 3 | B |
| Hymenoptera | Halictidae | *Lasioglossum coactus* (Cresson) | This study, (Michener 2007) | 203 | All |
| Hymenoptera | Halictidae | *Lasioglossum connexum* (Cresson) | This study, (Michener 2007) | 25 | A, B, D |
| Hymenoptera | Halictidae | *Lasioglossum coreopsis* (Robertson) | This study, (Michener 2007) | 47 | A, D |
| Hymenoptera | Halictidae | *Lasioglossum disparile* (Cresson) | This study, (Michener 2007) | 41 | A, C, D |
| Hymenoptera | Halictidae | *Lasioglossum fedorense* (Crawford) | This study, (Michener 2007) | 3 | D |
| Hymenoptera | Halictidae | *Lasioglossum hitchensi* Gibbs | (Michener 2007) | 11 | A-C |
| Hymenoptera | Halictidae | *Lasioglossum hudsoniellum* (Cockerell) | (Michener 2007) | 8 | B |
| Hymenoptera | Halictidae | *Lasioglossum ellisiae* (Sandhouse) | (Michener 2007) | 2 | A, D |
| Hymenoptera | Halictidae | *Lasioglossum illinoense* (Robertson) | (Michener 2007) | 15 | A |
| Hymenoptera | Halictidae | *Lasioglossum lustrans* (Cockerell) | This study, (Michener 2007) | 2 | D |
| Hymenoptera | Halictidae | *Lasioglossum pectorale* (Smith) | (Michener 2007) | 2 | D |
| Hymenoptera | Halictidae | *Lasioglossum* (Dialictus) sp. OK-1 | This study, (Michener 2007) | 5 | B |
| Hymenoptera | Halictidae | *Lasioglossum* (Dialictus) sp. TX-24 | This study, (Michener 2007) | 21 | B, D |
| Hymenoptera | Halictidae | *Lasioglossum* (Dialictus) sp. TX-27 | (Michener 2007) | 1 | D |
| Hymenoptera | Halictidae | *Lasioglossum* (Dialictus) sp. TX-3 | This study, (Michener 2007) | 139 | All |
| Hymenoptera | Halictidae | *Lasioglossum* (Dialictus) sp. TX-6 | (Michener 2007) | 3 | C |

| Order | Family | Scientific name/morphospecies ID | Visits flowers | Abundance | Periods present |
|---|---|---|---|---|---|
| Hymenoptera | Halictidae | *Lasioglossum tegulare* (Robertson) | This study, (Michener 2007) | 67 | All |
| Hymenoptera | Halictidae | *Lasioglossum* sp. | This study, (Michener 2007) | 2 | A |
| Hymenoptera | Halictidae | *Sphecodes* sp. TX-5 | (Michener 2007) | 1 | A |
| Hymenoptera | Halictidae | *Sphecodes mandibularis* Cresson | (Michener 2007) | 1 | D |
| Hymenoptera | Ichneumonidae | *Enicospilus* sp. | (Gauld & Wahl 2024) | 1 | B |
| Hymenoptera | Ichneumonidae | sp. 1 | (Pascarella et al. 2001) | 1 | C |
| Hymenoptera | Megachilidae | *Anthidiellum notatum* (Latreille) | This study, (Michener 2007) | 2 | B |
| Hymenoptera | Megachilidae | *Dianthidium curvatum* (Smith) | This study, (Michener 2007) | 7 | B, C |
| Hymenoptera | Megachilidae | *Dianthidium texanum* (Cresson) | This study, (Michener 2007) | 5 | A, B |
| Hymenoptera | Megachilidae | *Heriades carinata* Cresson | This study, (Michener 2007) | 2 | C |
| Hymenoptera | Megachilidae | *Megachile addenda* Cresson | This study, (Michener 2007) | 1 | D |
| Hymenoptera | Megachilidae | *Megachile albitarsis* (Cresson) | This study, (Michener 2007) | 5 | C, D |
| Hymenoptera | Megachilidae | *Megachile brevis* Say | This study, (Michener 2007) | 38 | All |
| Hymenoptera | Megachilidae | *Megachile comata* Cresson | This study, (Michener 2007) | 3 | B, C |
| Hymenoptera | Megachilidae | *Megachile fortis* Cresson | This study, (Michener 2007) | 1 | B |
| Hymenoptera | Megachilidae | *Megachile inimica* Cresson | This study, (Michener 2007) | 2 | C |

| **Order** | **Family** | **Scientific name/morphospecies ID** | **Visits flowers** | **Abundance** | **Periods present** |
|---|---|---|---|---|---|
| Hymenoptera | Megachilidae | *Megachile mendica* Cresson | This study, (Michener 2007) | 4 | C |
| Hymenoptera | Megachilidae | *Megachile montivaga* Cresson | This study, (Michener 2007) | 10 | A, C |
| Hymenoptera | Megachilidae | *Megachile parallela* Smith | This study, (Michener 2007) | 15 | All |
| Hymenoptera | Megachilidae | *Megachile policaris* Say | This study, (Michener 2007) | 9 | A-C |
| Hymenoptera | Megachilidae | *Megachile pugnata* Say | This study, (Michener 2007) | 1 | C |
| Hymenoptera | Megachilidae | *Osmia chalybea* Smith | This study, (Michener 2007) | 3 | C |
| Hymenoptera | Megachilidae | *Osmia subfasciata* Cresson | This study, (Michener 2007) | 8 | A |
| Hymenoptera | Mymaridae | sp. | (Jervis et al. 1993) | 1 | A |
| Hymenoptera | Perilampidae | *Perilampus* sp. A | This study, (Johnson & Triplehorn 2004) | 1 | D |
| Hymenoptera | Pompilidae | *Ageniella accepta* (Cresson) | (Johnson & Triplehorn 2004) | 3 | B, C |
| Hymenoptera | Pompilidae | *Ageniella arcuate* (Banks) | (Johnson & Triplehorn 2004) | 6 | A, C |
| Hymenoptera | Pompilidae | *Ageniella placita* (Banks) | (Johnson & Triplehorn 2004) | 4 | A-C |
| Hymenoptera | Pompilidae | *Ageniella* sp. 5 | (Johnson & Triplehorn 2004) | 1 | A |
| Hymenoptera | Pompilidae | *Anoplius nigritus* (Dahlbom) | (Johnson & Triplehorn 2004) | 4 | A |
| Hymenoptera | Pompilidae | *Anoplius* sp. 3 | This study, (Johnson & Triplehorn 2004) | 5 | D |

| Order | Family | Scientific name/morphospecies ID | Visits flowers | Abundance | Periods present |
|---|---|---|---|---|---|
| Hymenoptera | Pompilidae | *Anoplius* sp. 6 | (Johnson & Triplehorn 2004) | 1 | D |
| Hymenoptera | Pompilidae | *Aporinellus yucatanensis* (Cameron) | (Johnson & Triplehorn 2004) | 10 | A, D |
| Hymenoptera | Pompilidae | *Arachnophroctonus* sp. 4 | (Johnson & Triplehorn 2004) | 1 | D |
| Hymenoptera | Pompilidae | *Entypus* sp. 2 | This study, (Pascarella et al. 2001; Johnson & Triplehorn 2004) | 1 | A |
| Hymenoptera | Pompilidae | *Notiochares* sp. 1 | This study, (Johnson & Triplehorn 2004) | 1 | A |
| Hymenoptera | Pteromalidae | *Pteromalus* sp. A | (Jervis et al. 1993) | 1 | D |
| Hymenoptera | Pteromalidae | sp. d | (Jervis et al. 1993) | 1 | A |
| Hymenoptera | Pteromalidae | sp. f | (Jervis et al. 1993) | 1 | C |
| Hymenoptera | Pteromalinae | sp. a | (Jervis et al. 1993) | 1 | A |
| Hymenoptera | Sapygidae | *Sapyga louisi* Krombein | This study | 1 | C |
| Hymenoptera | Scelionidae | sp. 1 | (Orr 1988) | 1 | A |
| Hymenoptera | Scelionidae | sp. 2 | (Orr 1988) | 1 | A |
| Hymenoptera | Scelionidae | sp. g | (Orr 1988) | 2 | |
| Hymenoptera | Scoliidae | *Colpa octomaculata* (Say) | This study, (Johnson & Triplehorn 2004) | 1 | B |
| Hymenoptera | Scoliidae | *Dielis plumipes* (Drury) | This study, (Johnson & Triplehorn 2004) | 1 | C |
| Hymenoptera | Scoliidae | *Scolia dubia* Say | This study, (Johnson & Triplehorn 2004) | 1 | A |
| Hymenoptera | Sphecidae | *Ammophila* sp. 1 | (Evans 1970) | 1 | B |
| Hymenoptera | Sphecidae | *Sceliphron caementarium* (Drury) | This study | 2 | C |

| Order | Family | Scientific name/morphospecies ID | Visits flowers | Abundance | Periods present |
|---|---|---|---|---|---|
| Hymenoptera | Tiphiidae | *Myzinum maculate* (Fabricius) | This study, (Tooker & Hanks 2000) | 4 | A |
| Hymenoptera | Tiphiidae | *Myzinum quinquecinctum* (Fabricius) | This study, (Lavigne 1976; Tooker & Hanks 2000) | 10 | D |
| Hymenoptera | Tiphiidae | sp. A | This study | 1 | C |
| Hymenoptera | Trichogrammatidae | sp. | (Jervis et al. 1993) | 1 | B |
| Hymenoptera | Vespidae | *Euodynerus annulatus* (Say) | This study, (Lavigne 1976) | 1 | C |
| Hymenoptera | Vespidae | *Euodynerus pratensis* (de Saussure) | This study | 1 | A |
| Hymenoptera | Vespidae | *Parancistrocerus fulvipes* (de Saussure) | This study | 2 | C |
| Hymenoptera | Vespidae | *Parancistrocerus* sp. A | This study, (Wiesenborn 2017; Elmquist et al. 2022) | 1 | A |
| Hymenoptera | Vespidae | *Parancistrocerus* sp. B | This study, (Wiesenborn 2017; Elmquist et al. 2022) | 1 | A |
| Hymenoptera | Vespidae | *Polistes bellicosus* Cresson | This study, (Pascarella et al. 2001) | 13 | A, B |
| Hymenoptera | Vespidae | *Polistes carolina* (Linnaeus) | This study | 2 | A |
| Hymenoptera | Vespidae | *Polistes dorsalis* (Fabricius) | This study | 7 | A, B |
| Hymenoptera | Vespidae | *Polistes exclamans* Viereck | | 1 | A |
| Hymenoptera | Vespidae | *Polistes fuscatus* (Fabricius) | This study, (Lavigne 1976; Pascarella et al. 2001) | 2 | C, D |
| Hymenoptera | Vespidae | *Stenodynerus propinquus* (de Saussure) | This study, (Pascarella et al. 2001) | 3 | A |
| Hymenoptera | | sp. | | 1 | A |

| Order | Family | Scientific name/morphospecies ID | Visits flowers | Abundance | Periods present |
|---|---|---|---|---|---|
| Lepidoptera | Hesperiidae | *Amblyscirtes eos* (Edwards) | (Scott 1986) | 1 | B |
| Lepidoptera | Hesperiidae | *Amblyscirtes texanae* Bell | This study, (Scott 1986) | 1 | B |
| Lepidoptera | Hesperiidae | *Atalopedes campestris* (Boisduval) | This study, (Scott 1986) | 2 | B, C |
| Lepidoptera | Hesperiidae | *Burnsius communis* (Grote) | This study, (Scott 1986) | 2 | B |
| Lepidoptera | Hesperiidae | *Erynnis funeralis* (Scudder & Burgess) | This study, (Scott 1986) | 1 | A |
| Lepidoptera | Hesperiidae | *Lerema accius* (Smith) | This study, (Scott 1986) | 2 | D |
| Lepidoptera | Hesperiidae | *Lerodea eufala* (Edwards) | (Scott 1986) | 5 | A, B, D |
| Lepidoptera | Hesperiidae | *Polites origenes* (Fabricius) | (Scott 1986) | 1 | D |
| Lepidoptera | Lycaenidae | *Echinargus isola* (Reakirt) | This study, (Scott 1986) | 12 | A-C |
| Lepidoptera | Lycaenidae | *Strymon melinus* (Hübner) | This study, (Scott 1986) | 10 | B-D |
| Lepidoptera | Noctuidae | *Caradrina montana* (Bremer) | This study | 1 | C |
| Lepidoptera | Noctuidae | *Ponometia erastrioides* (Guenée) | This study | 1 | D |
| Lepidoptera | Nymphalidae | *Cercyonis pegala* (Fabricius) | This study, (Scott 1986) | 1 | B |
| Lepidoptera | Nymphalidae | *Danaus gilippus* (Cramer) | This study, (Scott 1986) | 1 | C |
| Lepidoptera | Nymphalidae | *Euptoieta claudia* (Cramer) | This study, (Scott 1986) | 1 | A |
| Lepidoptera | Nymphalidae | *Junonia coenia* Hübner | This study, (Scott 1986) | 2 | A, D |
| Lepidoptera | Nymphalidae | *Phyciodes tharos* (Drury) | This study, (Scott 1986) | 2 | C |
| Lepidoptera | Nymphalidae | *Vanessa cardui* (Linnaeus) | This study, (Scott 1986) | 1 | D |
| Lepidoptera | Nymphalidae | *Vanessa virginiensis* (Drury) | This study, (Scott 1986) | 2 | A, D |
| Lepidoptera | Pieridae | *Abaeis nicippe* (Cramer) | This study, (Scott 1986) | 8 | B |
| Lepidoptera | Pieridae | *Nathalis iole* Boisduval | This study, (Lavigne 1976; Scott 1986) | 7 | B, D |
| Lepidoptera | Pieridae | *Pontia protodice* (Boisduval & Leconte) | This study, (Scott 1986) | 2 | D |
| Lepidoptera | Pieridae | *Pyrisitia lisa* (Boisduval & Leconte) | This study, (Scott 1986) | 14 | A |
| Lepidoptera | Pterophoridae | *Adaina montanus* (Walshingham) | This study | 1 | A |

**Table S6: Coefficients and test statistics for best-fit models investigating effects of local and landscape habitat on Coleoptera abundance. ΔAICc indicates the difference between a given model's AICc value and that of the model with the smallest AICc. Shading separates models.**

| Landscape measure – model ranking | Model term | Coefficient | Coefficient standard error | $\chi^2$ | df | *P*-value | ΔAICc |
|---|---|---|---|---|---|---|---|
| Grassland – best 1 | Intercept | 2.41 | 0.28 | | | | 0 |
| Grassland – best 1 | Flower richness | -0.31 | 0.15 | 4.34 | 1 | 0.04 | |
| Grassland – best 2 | Intercept | 1.81 | 0.63 | | | | 0.87 |
| Grassland – best 2 | Landscape-scale grassland cover | 1.09 | 1.00 | 1.25 | 1 | 0.27 | |
| Grassland – best 2 | Flower richness | -0.32 | 0.15 | 4.99 | 1 | 0.03 | |
| Grassland – best 3 | Intercept | 1.82 | 0.61 | | | | 1.01 |
| Grassland – best 3 | Landscape-scale grassland cover | 1.12 | 0.97 | 3.47 | 2 | 0.18 | |
| Grassland – best 3 | Flower richness | -0.86 | 0.37 | 7.10 | 1 | 0.03 | |
| Grassland – best 3 | Grassland : Flower richness | 1.11 | 0.71 | 2.50 | 1 | 0.11 | |
| Woodland – best 1 | Intercept | 2.41 | 0.28 | | | | 0 |
| Woodland – best 1 | Flower richness | -0.31 | 0.15 | 4.34 | 1 | 0.04 | |
| Woodland – best 2 | Intercept | 1.94 | 0.35 | | | | 0.50 |

| **Landscape measure – model ranking** | **Model term** | **Coefficient** | **Coefficient standard error** | $\chi^2$ | **df** | ***P*-value** | **ΔAICc** |
|---|---|---|---|---|---|---|---|
| **Woodland – best 2** | **Landscape-scale woodland cover** | **1.64** | **0.98** | **2.01** | **1** | **0.16** | |
| **Woodland – best 2** | **Flower richness** | **-0.31** | **0.15** | **4.26** | **1** | **0.04** | |
| **Woodland – best 3** | **Intercept** | **2.44** | **0.28** | | | | **1.47** |

**Table S7: Coefficients and test statistics for best-fit models investigating effects of local and landscape habitat on Coleoptera richness. ΔAICc indicates the difference between a given model's AICc value and that of the model with the smallest AICc. Shading separates models.**

| Landscape measure – model ranking | Model term | Coefficient | Coefficient standard error | $\chi^2$ | df | *P*-value | ΔAICc |
|---|---|---|---|---|---|---|---|
| Grassland – best 1 | Intercept | 4.52 | 0.67 | | | | 0 |
| Woodland – best 1 | Intercept | 3.11 | 0.76 | | | | 0 |
| Grassland – best 1 | Landscape-scale grassland cover | 4.57 | 1.95 | 4.85 | 1 | 0.03 | |

**Table S8: Coefficients and test statistics for best-fit models investigating effects of local and landscape habitat on Coleoptera evenness. ΔAICc indicates the difference between a given model's AICc value and that of the model with the smallest AICc. Shading separates models.**

| Landscape measure – model ranking | Model term | Coefficient | Coefficient standard error | $\chi^2$ | df | *P*-value | ΔAICc |
|---|---|---|---|---|---|---|---|
| Grassland – best 1 | Intercept | 0.84 | 0.11 | | | | 0 |
| Grassland – best 1 | Landscape-scale grassland cover | -0.16 | 0.16 | 7.75 | 2 | 0.02 | |
| Grassland – best 1 | Flower richness | 0.26 | 0.06 | 21.31 | 2 | <0.0001 | |
| Grassland – best 1 | Invasive perennial grass cover | -0.13 | 0.06 | 5.56 | 1 | 0.02 | |
| Grassland – best 1 | Grassland : Flower richness | -0.32 | 0.12 | 7.18 | 1 | 0.01 | |
| Grassland – best 2 | Intercept | 0.72 | 0.06 | | | | 1.10 |
| Grassland – best 2 | Flower richness | 0.10 | 0.03 | 12.18 | 1 | 0.0005 | |
| Grassland – best 3 | Intercept | 0.75 | 0.06 | | | | 1.67 |
| Grassland – best 3 | Flower richness | 0.10 | 0.03 | 13.59 | 1 | 0.0002 | |
| Grassland – best 3 | Invasive perennial grass cover | -0.09 | 0.06 | 2.17 | 1 | 0.14 | |
| Woodland – best 1 | Intercept | 0.85 | 0.07 | | | | 0 |
| Woodland – best 1 | Landscape-scale woodland cover | -0.42 | 0.16 | 4.75 | 1 | 0.03 | |

| Landscape measure – model ranking | Model term | Coefficient | Coefficient standard error | $\chi^2$ | df | *P*-value | ΔAICc |
|---|---|---|---|---|---|---|---|
| Woodland – best 1 | Flower richness | 0.10 | 0.03 | 11.75 | 1 | 0.0006 | |
| Woodland – best 2 | Intercept | 0.83 | 0.06 | | | | 0.89 |
| Woodland – best 2 | Landscape-scale woodland cover | -0.39 | 0.16 | 6.79 | 2 | 0.03 | |
| Woodland – best 2 | Flower richness | 0.04 | 0.05 | 13.79 | 2 | 0.001 | |
| Woodland – best 2 | Woodland: Flower richness | 0.14 | 0.10 | 2.04 | 1 | 0.15 | |
| Woodland – best 3 | Intercept | 0.86 | 0.07 | | | | 1.89 |
| Woodland – best 3 | Landscape-scale woodland cover | -0.38 | 0.16 | 3.63 | 1 | 0.057 | |
| Woodland – best 3 | Flower richness | 0.10 | 0.03 | 12.51 | 1 | 0.0004 | |
| Woodland – best 3 | Invasive perennial grass cover | -0.08 | 0.07 | 1.04 | 1 | 0.31 | |

**Table S9: Coefficients and test statistics for best-fit models investigating effects of local and landscape habitat on Diptera abundance. ΔAICc indicates the difference between a given model's AICc value and that of the model with the smallest AICc. Shading separates models.**

| Landscape measure – model ranking | Model term | Coefficient | Coefficient standard error | $\chi^2$ | df | *P*-value | ΔAICc |
|---|---|---|---|---|---|---|---|
| Grassland – best 1 | Intercept | 3.12 | 0.39 | | | | 0 |
| Grassland – best 1 | Landscape-scale grassland cover | 0.99 | 0.66 | 2.44 | 1 | 0.12 | |
| Grassland – best 1 | Bare ground cover | -9.03 | 3.54 | 4.57 | 1 | 0.03 | |
| Grassland – best 2 | Intercept | 3.65 | 0.16 | | | | 0.04 |
| Grassland – best 2 | Bare ground cover | -7.70 | 3.59 | 4.04 | 1 | 0.04 | |
| Grassland – best 3 | Intercept | 3.54 | 0.16 | | | | 1.96 |
| Woodland – best 1 | Intercept | 3.65 | 0.16 | | | | 0 |
| Woodland – best 1 | Bare ground cover | -7.70 | 3.59 | 4.04 | 1 | 0.04 | |
| Woodland – best 2 | Intercept | 3.90 | 0.23 | | | | 0.98 |
| Woodland – best 2 | Landscape-scale woodland cover | -0.78 | 0.59 | 1.63 | 1 | 0.20 | |
| Woodland – best 2 | Bare ground cover | -7.39 | 3.61 | 3.72 | 1 | 0.05 | |
| Woodland – best 3 | Intercept | 3.54 | 0.16 | | | | 1.96 |

**Table S10: Coefficients and test statistics for best-fit models investigating effects of local and landscape habitat on Diptera richness. ΔAICc indicates the difference between a given model's AICc value and that of the model with the smallest AICc. Shading separates models.**

| Landscape measure – model ranking | Model term | Coefficient | Coefficient standard error | $\chi^2$ | df | *P*-value | ΔAICc |
|---|---|---|---|---|---|---|---|
| Grassland – best 1 | Intercept | 2.03 | 0.11 | | | | 0 |
| Grassland – best 2 | Intercept | 2.11 | 0.12 | | | | 0.66 |
| Grassland – best 2 | Invasive perennial grass cover | -0.25 | 0.19 | 1.65 | 1 | 0.20 | |
| Grassland – best 3 | Intercept | 1.77 | 0.24 | | | | 0.97 |
| Grassland – best 3 | Landscape-scale grassland cover | 0.49 | 0.40 | 1.35 | 1 | 0.25 | |
| Woodland – best 1 | Intercept | 2.29 | 0.14 | | | | 0 |
| Woodland – best 1 | Landscape-scale woodland cover | -0.85 | 0.37 | 3.96 | 1 | 0.047 | |
| Woodland – best 2 | Intercept | 2.37 | 0.15 | | | | 0.88 |
| Woodland – best 2 | Landscape-scale woodland cover | -0.85 | 0.38 | 3.86 | 1 | 0.049 | |
| Woodland – best 2 | Invasive perennial grass cover | -0.23 | 0.19 | 1.56 | 1 | 0.21 | |
| Woodland – best 3 | Intercept | 2.03 | 0.11 | | | | 1.64 |

**Table S11: Coefficients and test statistics for best-fit models investigating effects of local and landscape habitat on Diptera evenness. ΔAICc indicates the difference between a given model's AICc value and that of the model with the smallest AICc. Shading separates models.**

| Landscape measure – model ranking | Model term | Coefficient | Coefficient standard error | $\chi^2$ | df | *P*-value | ΔAICc |
|---|---|---|---|---|---|---|---|
| Grassland or woodland – best 1 | Intercept | -0.68 | 0.06 | | | | 0 |
| Grassland or woodland – best 2 | Intercept | -0.61 | 0.08 | | | | 0.97 |
| Grassland or woodland – best 2 | Invasive perennial grass cover | -0.20 | 0.16 | 1.48 | 1 | 0.23 | |
| Grassland or woodland – best 3 | Intercept | -0.68 | 0.06 | | | | 1.98 |
| Grassland or woodland – best 3 | Flower richness | 0.04 | 0.06 | 0.47 | 1 | 0.49 | |

**Table S12: Coefficients and test statistics for best-fit models investigating effects of local and landscape habitat on Hymenoptera abundance. ΔAICc indicates the difference between a given model's AICc value and that of the model with the smallest AICc. Shading separates models.**

| Landscape measure – model ranking | Model term | Coefficient | Coefficient standard error | $\chi^2$ | df | *P*-value | ΔAICc |
|---|---|---|---|---|---|---|---|
| Grassland – best 1 | Intercept | 3.51 | 0.15 | | | | 0 |
| Grassland – best 1 | Flower richness | 0.28 | 0.11 | 5.77 | 1 | 0.02 | |
| Grassland – best 1 | Bare ground cover | 6.34 | 2.71 | 6.01 | 1 | 0.02 | |
| Grassland – best 2 | Intercept | 3.66 | 0.18 | | | | 0.24 |
| Grassland – best 2 | Flower richness | 0.29 | 0.11 | 6.36 | 1 | 0.01 | |
| Grassland – best 2 | Bare ground cover | 5.91 | 2.60 | 5.60 | 1 | 0.02 | |
| Grassland – best 2 | Invasive perennial grass cover | -0.48 | 0.31 | 2.47 | 1 | 0.12 | |
| Grassland – best 3 | Intercept | 3.22 | 0.31 | | | | 1.74 |
| Grassland – best 3 | Landscape-scale grassland cover | 0.58 | 0.55 | 1.04 | 1 | 0.31 | |
| Grassland – best 3 | Flower richness | 0.29 | 0.11 | 6.30 | 1 | 0.01 | |
| Grassland – best 3 | Bare ground cover | 5.91 | 2.75 | 4.96 | 1 | 0.03 | |
| Woodland – best 1 | Intercept | 3.83 | 0.19 | | | | 0 |

| Landscape measure – model ranking | Model term | Coefficient | Coefficient standard error | $\chi^2$ | df | *P*-value | ΔAICc |
|---|---|---|---|---|---|---|---|
| Woodland – best 1 | Landscape-scale woodland cover | -1.00 | 0.46 | 3.44 | 1 | 0.06 | |
| Woodland – best 1 | Flower richness | 0.30 | 0.11 | 2.73 | 1 | 0.007 | |
| Woodland – best 1 | Bare ground cover | 6.74 | 2.78 | 6.47 | 1 | 0.01 | |
| Woodland – best 2 | Intercept | 3.51 | 0.15 | | | | 0.58 |
| Woodland – best 2 | Flower richness | 0.28 | 0.11 | 5.77 | 1 | 0.02 | |
| Woodland – best 2 | Bare ground cover | 6.34 | 2.71 | 6.01 | 1 | 0.01 | |
| Woodland – best 3 | Intercept | 3.97 | 0.22 | | | | 0.71 |
| Woodland – best 3 | Landscape-scale woodland cover | -1.00 | 0.50 | 3.18 | 1 | 0.07 | |
| Woodland – best 3 | Flower richness | 0.31 | 0.11 | 7.70 | 1 | 0.006 | |
| Woodland – best 3 | Bare ground cover | 6.13 | 2.67 | 5.72 | 1 | 0.02 | |
| Woodland – best 3 | Invasive perennial grass cover | -0.45 | 0.30 | 2.25 | 1 | 0.13 | |
| Woodland – best 4 | Intercept | 3.66 | 0.18 | | | | 1.06 |
| Woodland – best 4 | Flower richness | 0.29 | 0.11 | 6.36 | 1 | 0.01 | |
| Woodland – best 4 | Bare ground cover | 5.81 | 2.60 | 5.60 | 1 | 0.02 | |

| Landscape measure – model ranking | Model term | Coefficient | Coefficient standard error | $\chi^2$ | df | *P*-value | ΔAICc |
|---|---|---|---|---|---|---|---|
| **Woodland – best 4** | **Invasive perennial grass cover** | **-0.48** | **0.31** | **2.50** | **1** | **0.12** | |
| **Woodland – best 5** | **Intercept** | **3.86** | **0.18** | | | | **1.61** |
| **Woodland – best 5** | **Landscape-scale woodland cover** | **-1.08** | **0.44** | **4.66** | **2** | **0.097** | |
| **Woodland – best 5** | **Flower richness** | **0.47** | **0.17** | **8.48** | **2** | **0.01** | |
| **Woodland – best 5** | **Bare ground cover** | **7.04** | **2.83** | **6.85** | **1** | **0.009** | |
| **Woodland – best 5** | **Woodland : Flower richness** | **-0.51** | **0.45** | **1.26** | **1** | **0.26** | |

**Table S13: Coefficients and test statistics for best-fit models investigating effects of local and landscape habitat on Hymenoptera richness. ΔAICc indicates the difference between a given model's AICc value and that of the model with the smallest AICc. Shading separates models.**

| Landscape measure – model ranking | Model term | Coefficient | Coefficient standard error | $\chi^2$ | df | *P*-value | ΔAICc |
|---|---|---|---|---|---|---|---|
| Grassland or woodland – best 1 | Intercept | 12.48 | 0.89 | | | | 0 |
| Grassland or woodland – best 1 | Flower richness | 1.70 | 0.70 | 5.72 | 1 | 0.02 | |
| Grassland or woodland – best 2 | Intercept | 13.30 | 1.10 | | | | 0.83 |
| Grassland or woodland – best 2 | Flower richness | 1.79 | 0.70 | 6.34 | 1 | 0.01 | |
| Grassland or woodland – best 2 | Invasive perennial grass cover | -2.58 | 1.95 | 1.74 | 1 | 0.19 | |

**Table S14: Coefficients and test statistics for best-fit models investigating effects of local and landscape habitat on Hymenoptera evenness. ΔAICc indicates the difference between a given model's AICc value and that of the model with the smallest AICc. Shading separates models.**

| Landscape measure – model ranking | Model term | Coefficient | Coefficient standard error | $\chi^2$ | df | *P*-value | ΔAICc |
|---|---|---|---|---|---|---|---|
| Grassland – best 1 | Intercept | 0.68 | 0.04 | | | | 0 |
| Grassland – best 1 | Bare ground cover | -1.81 | 0.71 | 6.13 | 1 | 0.01 | |
| Grassland – best 2 | Intercept | 0.64 | 0.05 | | | | 0.31 |
| Grassland – best 2 | Bare ground cover | -1.69 | 0.69 | 5.92 | 1 | 0.02 | |
| Grassland – best 2 | Invasive perennial grass cover | 0.11 | 0.07 | 2.26 | 1 | 0.13 | |
| Grassland – best 3 | Intercept | 0.68 | 0.04 | | | | 0.83 |
| Grassland – best 3 | Flower richness | -0.03 | 0.03 | 1.74 | 1 | 0.19 | |
| Grassland – best 3 | Bare ground cover | -1.79 | 0.72 | 6.14 | 1 | 0.01 | |
| Grassland – best 4 | Intercept | 0.64 | 0.05 | | | | 0.94 |
| Grassland – best 4 | Flower richness | -0.04 | 0.03 | 2.08 | 1 | 0.15 | |
| Grassland – best 4 | Bare ground cover | -1.67 | 0.70 | 5.88 | 1 | 0.02 | |
| Grassland – best 4 | Invasive perennial grass cover | 0.12 | 0.07 | 2.60 | 1 | 0.11 | |

| Landscape measure – model ranking | Model term | Coefficient | Coefficient standard error | $\chi^2$ | df | *P*-value | ΔAICc |
|---|---|---|---|---|---|---|---|
| Grassland – best 5 | Intercept | 0.74 | 0.09 | | | | 1.88 |
| Grassland – best 5 | Landscape-scale grassland cover | -0.13 | 0.16 | 0.69 | 1 | 0.41 | |
| Grassland – best 5 | Bare ground cover | -1.71 | 0.73 | 5.55 | 1 | 0.02 | |
| Woodland – best 1 | Intercept | 0.68 | 0.04 | | | | 0 |
| Woodland – best 1 | Bare ground cover | -1.81 | 0.71 | 6.13 | 1 | 0.01 | |
| Woodland – best 2 | Intercept | 0.61 | 0.06 | | | | 0.24 |
| Woodland – best 2 | Landscape-scale woodland cover | 0.22 | 0.15 | 2.33 | 1 | 0.13 | |
| Woodland – best 2 | Bare ground cover | -1.80 | 0.71 | 6.29 | 1 | 0.01 | |
| Woodland – best 3 | Intercept | 0.64 | 0.05 | | | | 0.31 |
| Woodland – best 3 | Bare ground cover | -1.69 | 0.69 | 5.92 | 1 | 0.02 | |
| Woodland – best 3 | Invasive perennial grass cover | 0.11 | 0.07 | 2.26 | 1 | 0.13 | |
| Woodland – best 4 | Intercept | 0.61 | 0.06 | | | | 0.63 |
| Woodland – best 4 | Landscape-scale woodland cover | 0.23 | 0.14 | 2.91 | 1 | 0.09 | |
| Woodland – best 4 | Flower richness | -0.04 | 0.03 | 2.32 | 1 | 0.13 | |

| Landscape measure – model ranking | Model term | Coefficient | Coefficient standard error | $\chi^2$ | df | *P*-value | ΔAICc |
|---|---|---|---|---|---|---|---|
| Woodland – best 4 | Bare ground cover | -1.80 | 0.71 | 6.47 | 1 | 0.01 | |
| Woodland – best 5 | Intercept | 0.57 | 0.07 | | | | 0.73 |
| Woodland – best 5 | Landscape-scale woodland cover | 0.22 | 0.16 | 2.28 | 1 | 0.13 | |
| Woodland – best 5 | Bare ground cover | -1.66 | 0.69 | 5.92 | 1 | 0.02 | |
| Woodland – best 5 | Invasive perennial grass cover | 0.12 | 0.07 | 2.21 | 1 | 0.14 | |
| Woodland – best 6 | Intercept | 0.56 | 0.06 | | | | 0.83 |
| Woodland – best 6 | Landscape-scale woodland cover | 0.23 | 0.15 | 2.90 | 1 | 0.09 | |
| Woodland – best 6 | Flower richness | -0.04 | 0.03 | 2.70 | 1 | 0.10 | |
| Woodland – best 6 | Bare ground cover | -1.65 | 0.69 | 6.07 | 1 | 0.01 | |
| Woodland – best 6 | Invasive perennial grass cover | 0.12 | 0.07 | 2.59 | 1 | 0.11 | |
| Woodland – best 7 | Intercept | 0.68 | 0.04 | | | | 0.91 |
| Woodland – best 7 | Flower richness | -0.03 | 0.03 | 1.74 | 1 | 0.19 | |
| Woodland – best 7 | Bare ground cover | -1.79 | 0.72 | 6.14 | 1 | 0.01 | |
| Woodland – best 8 | Intercept | 0.64 | 0.05 | | | | 0.94 |

| Landscape measure – model ranking | Model term | Coefficient | Coefficient standard error | $\chi^2$ | df | *P*-value | ΔAICc |
|---|---|---|---|---|---|---|---|
| **Woodland – best 8** | **Flower richness** | **-0.04** | **0.03** | **2.08** | **1** | **0.15** | |
| **Woodland – best 8** | **Bare ground cover** | **-1.67** | **0.70** | **5.88** | **1** | **0.02** | |
| **Woodland – best 8** | **Invasive perennial grass cover** | **0.12** | **0.07** | **2.60** | **1** | **0.11** | |

**Table S15: Coefficients and test statistics for best-fit models investigating effects of local and landscape habitat on Lepidoptera presence. ΔAICc indicates the difference between a given model's AICc value and that of the model with the smallest AICc. Shading separates models.**

| Landscape measure – model ranking | Model term | Coefficient | Coefficient standard error | $\chi^2$ | df | *P*-value | ΔAICc |
|---|---|---|---|---|---|---|---|
| Grassland – best 1 | Intercept | 0.81 | 0.82 | | | | 0 |
| Grassland – best 1 | Bare ground cover | 79.52 | 50.75 | 7.29 | 1 | 0.007 | |
| Grassland – best 2 | Intercept | -1.09 | 1.73 | | | | 1.29 |
| Grassland – best 2 | Landscape-scale grassland cover | 3.22 | 2.86 | 1.15 | 1 | 0.28 | |
| Grassland – best 2 | Bare ground cover | 91.90 | 55.07 | 7.65 | 1 | 0.006 | |
| Woodland – best 1 | Intercept | 0.81 | 0.82 | | | | 0 |
| Woodland – best 1 | Bare ground cover | 79.52 | 50.75 | 7.29 | 1 | 0.007 | |

**Table S16: Coefficients and test statistics for PERMANOVAs investigating effects of local and landscape habitat on Diptera community composition. Shading separates models.**

| Landscape measure | Model term | $r^2$ | F | df | *P*-value |
|---|---|---|---|---|---|
| Grassland | Landscape-scale grassland cover | 0.03 | 1.35 | 1 | 0.21 |
| Grassland | Flower richness | 0.02 | 0.76 | 1 | 0.65 |
| Grassland | Bare ground cover | 0.06 | 2.56 | 1 | 0.01 |
| Grassland | Invasive perennial grass cover | 0.02 | 0.94 | 1 | 0.49 |
| Grassland | Grassland : Flower richness | 0.03 | 1.40 | 1 | 0.18 |
| Grassland | Residual | | | 37 | |
| Woodland | Landscape-scale woodland cover | 0.08 | 4.06 | 1 | 0.0009 |
| Woodland | Flower richness | 0.02 | 1.03 | 1 | 0.40 |
| Woodland | Bare ground cover | 0.07 | 3.49 | 1 | 0.002 |
| Woodland | Invasive perennial grass cover | 0.03 | 1.59 | 1 | 0.12 |
| Woodland | Woodland : Flower richness | 0.06 | 2.70 | 1 | 0.01 |
| Woodland | Residual | | | 37 | |

**Table S17: Coefficients and test statistics for PERMANOVAs investigating effects of local and landscape habitat on Hymenoptera community composition. Shading separates models.**

| Landscape measure | Model term | $r^2$ | F | df | *P*-value |
|---|---|---|---|---|---|
| Grassland | Landscape-scale grassland cover | 0.02 | 0.91 | 1 | 0.56 |
| Grassland | Flower richness | 0.03 | 1.27 | 1 | 0.21 |
| Grassland | Bare ground cover | 0.02 | 0.88 | 1 | 0.60 |
| Grassland | Invasive perennial grass cover | 0.03 | 1.23 | 1 | 0.24 |
| Grassland | Grassland : Flower richness | 0.02 | 0.91 | 1 | 0.56 |
| Grassland | Residual | | | 37 | |
| Woodland | Landscape-scale woodland cover | 0.03 | 1.51 | 1 | 0.09 |
| Woodland | Flower richness | 0.03 | 1.20 | 1 | 0.006 |
| Woodland | Bare ground cover | 0.03 | 1.30 | 1 | 0.19 |
| Woodland | Invasive perennial grass cover | 0.04 | 1.66 | 1 | 0.05 |
| Woodland | Woodland : Flower richness | 0.05 | 2.24 | 1 | 0.006 |
| Woodland | Residual | | | 37 | |

**Table S18: Coefficients and test statistics for PERMANOVAs investigating effects of local and landscape habitat on Coleoptera community composition. Shading separates models.**

| Landscape measure | Model term | $r^2$ | F | df | *P*-value |
|---|---|---|---|---|---|
| Grassland | Landscape-scale grassland cover | 0.01 | 0.31 | 1 | 0.92 |
| Grassland | Flower richness | 0.04 | 1.79 | 1 | 0.11 |
| Grassland | Bare ground cover | 0.04 | 1.60 | 1 | 0.15 |
| Grassland | Invasive perennial grass cover | 0.03 | 1.22 | 1 | 0.31 |
| Grassland | Grassland : Flower richness | 0.03 | 1.01 | 1 | 0.44 |
| Grassland | Residual | | | 33 | |
| Woodland | Landscape-scale woodland cover | 0.02 | 0.85 | 1 | 0.54 |
| Woodland | Flower richness | 0.05 | 1.95 | 1 | 0.09 |
| Woodland | Bare ground cover | 0.05 | 1.85 | 1 | 0.10 |
| Woodland | Invasive perennial grass cover | 0.03 | 1.20 | 1 | 0.32 |
| Woodland | Woodland : Flower richness | 0.01 | 0.52 | 1 | 0.78 |
| Woodland | Residual | | | 33 | |

**Table S19:** Test statistics for indicator species analyses investigating impacts of bare ground cover on Diptera community composition. “1” values indicate which habitat category each taxon is associated with.

| Landscape measure | Taxon | Less bare ground | More bare ground | Test statistic | *P*-value |
|---|---|---|---|---|---|
| Grassland | Asilidae *Mallophora orcina* | 1 | 0 | 0.22 | 0.4931 |
| Grassland | Asilidae *Ospriocerus* spp. | 0 | 1 | 0.28 | 0.1093 |
| Grassland | Asilidae *Psilocurus* spp. | 0 | 1 | 0.01 | 1.0000 |
| Grassland | Asteiidae *Leiomyza* spp. | 1 | 0 | 0.09 | 1.0000 |
| Grassland | Bombyliidae *Chrysanthrax cypris* | 1 | 0 | 0.05 | 1.0000 |
| Grassland | Bombyliidae *Chrysanthrax edititius* | 0 | 1 | 0.01 | 1.0000 |
| Grassland | Bombyliidae *Geron* spp. | 0 | 1 | 0.35 | 0.0352 |
| Grassland | Bombyliidae *Hemipenthes sinuosa* | 0 | 1 | 0.01 | 1.0000 |
| Grassland | Bombyliidae *Poecilanthrax lucifer* | 0 | 1 | 0.01 | 1.0000 |
| Grassland | Bombyliidae *Poecilognathus punctipennis* | 0 | 1 | 0.10 | 0.6073 |
| Grassland | Bombyliidae *Poecilognathus* sp. 1 | 1 | 0 | 0.05 | 1.0000 |
| Grassland | Bombyliidae *Rhynchanthrax* spp. | 0 | 1 | 0.01 | 1.0000 |
| Grassland | Bombyliidae *Systoechus solitus* | 1 | 0 | 0.27 | 0.2385 |
| Grassland | Bombyliidae *Villa lateralis* | 1 | 0 | 0.15 | 0.6038 |
| Grassland | Calliphoridae *Lucilia* spp. | 0 | 1 | 0.01 | 1.0000 |
| Grassland | Chloropidae *Apallates* spp. | 0 | 1 | 0.07 | 0.6918 |
| Grassland | Chloropidae *Chlorops* spp. | 1 | 0 | 0.05 | 1.0000 |
| Grassland | Chloropidae *Conioscinella* spp. | 1 | 0 | 0.09 | 0.7299 |
| Grassland | Chloropidae *Hippelates* spp. | 0 | 1 | 0.01 | 1.0000 |
| Grassland | Chloropidae *Incertella* spp. | 1 | 0 | 0.33 | 0.0577 |
| Grassland | Dolichopodidae *Asyndetus* spp. | 0 | 1 | 0.17 | 0.3082 |
| Grassland | Dolichopodidae *Chrysotus* spp. | 1 | 0 | 0.02 | 1.0000 |
| Grassland | Dolichopodidae *Condylostylus* spp. | 1 | 0 | 0.65 | 0.0001 |

| **Landscape measure** | **Taxon** | **Less bare ground** | **More bare ground** | **Test statistic** | ***P*-value** |
|---|---|---|---|---|---|
| Grassland | Fannidae *Fannia pusio* | 1 | 0 | 0.15 | 1.0000 |
| Grassland | Hybotidae Tachydromiinae spp. | 1 | 0 | 0.27 | 0.2374 |
| Grassland | Muscidae *Atherigona orientalis* | 1 | 0 | 0.05 | 1.0000 |
| Grassland | Muscidae *Caricea erythrocera* | 1 | 0 | 0.22 | 0.4842 |
| Grassland | Muscidae *Polietes* sp. 1 | 1 | 0 | 0.09 | 1.0000 |
| Grassland | Muscidae spp. | 0 | 1 | 0.01 | 1.0000 |
| Grassland | Phoridae Group 1 | 1 | 0 | 0.21 | 0.3483 |
| Grassland | Phoridae Group 2 | 1 | 0 | 0.18 | 0.3209 |
| Grassland | Pipunculidae *Tomosvaryella* spp. | 0 | 1 | 0.01 | 1.0000 |
| Grassland | Sarcophagidae *Argoravinia modesta* | 1 | 0 | 0.21 | 0.3409 |
| Grassland | Sarcophagidae *Bercaea cruentata* | 1 | 0 | 0.22 | 0.4972 |
| Grassland | Sarcophagidae *Comasarcophaga* spp. | 1 | 0 | 0.09 | 1.0000 |
| Grassland | Sarcophagidae *Helicobia rapax* | 1 | 0 | 0.43 | 0.0091 |
| Grassland | Sarcophagidae Miltogramminae spp. | 1 | 0 | 0.32 | 0.1089 |
| Grassland | Sarcophagidae *Ravinia* spp. | 1 | 0 | 0.22 | 0.2361 |
| Grassland | Sarcophagidae *Tylomyia* spp. | 1 | 0 | 0.09 | 1.0000 |
| Grassland | Sciaridae *Eugnoriste brevirostris* | 1 | 0 | 0.15 | 1.0000 |
| Grassland | Sciaridae spp. | 1 | 0 | 0.36 | 0.0494 |
| Grassland | Stratiomyidae *Nemotelus kansensis* | 0 | 1 | 0.01 | 1.0000 |
| Grassland | Syrphidae *Copestylum tamaulipanum* | 0 | 1 | 0.01 | 1.0000 |
| Grassland | Syrphidae *Palpada vinetorum* | 1 | 0 | 0.22 | 0.4880 |
| Grassland | Syrphidae *Toxomerus marginatus* | 0 | 1 | 0.10 | 0.6123 |
| Grassland | Tachinidae *Cylindromyia intermedia* | 0 | 1 | 0.08 | 0.6669 |
| Grassland | Tachinidae *Phasia* spp. | 1 | 0 | 0.22 | 0.4937 |
| Woodland | Asilidae *Mallophora orcina* | 1 | 0 | 0.22 | 0.4867 |

| Landscape measure | Taxon | Less bare ground | More bare ground | Test statistic | *P*-value |
|---|---|---|---|---|---|
| Woodland | Asilidae *Ospriocerus* spp. | 0 | 1 | 0.28 | 0.1057 |
| Woodland | Asilidae *Psilocurus* spp. | 0 | 1 | 0.01 | 1.0000 |
| Woodland | Asteiidae *Leiomyza* spp. | 1 | 0 | 0.09 | 1.0000 |
| Woodland | Bombyliidae *Chrysanthrax cypris* | 1 | 0 | 0.05 | 1.0000 |
| Woodland | Bombyliidae *Chrysanthrax edititius* | 0 | 1 | 0.01 | 1.0000 |
| Woodland | Bombyliidae *Geron* spp. | 0 | 1 | 0.35 | 0.0372 |
| Woodland | Bombyliidae *Hemipenthes sinuosa* | 0 | 1 | 0.01 | 1.0000 |
| Woodland | Bombyliidae *Poecilanthrax lucifer* | 0 | 1 | 0.01 | 1.0000 |
| Woodland | Bombyliidae *Poecilognathus punctipennis* | 0 | 1 | 0.10 | 0.6087 |
| Woodland | Bombyliidae *Poecilognathus* sp. 1 | 1 | 0 | 0.05 | 1.0000 |
| Woodland | Bombyliidae *Rhynchanthrax* spp. | 0 | 1 | 0.01 | 1.0000 |
| Woodland | Bombyliidae *Systoechus solitus* | 1 | 0 | 0.27 | 0.2296 |
| Woodland | Bombyliidae *Villa lateralis* | 1 | 0 | 0.15 | 0.5986 |
| Woodland | Calliphoridae *Lucilia* spp. | 0 | 1 | 0.01 | 1.0000 |
| Woodland | Chloropidae *Apallates* spp. | 0 | 1 | 0.07 | 0.6986 |
| Woodland | Chloropidae *Chlorops* spp. | 1 | 0 | 0.05 | 1.0000 |
| Woodland | Chloropidae *Conioscinella* spp. | 1 | 0 | 0.09 | 0.7388 |
| Woodland | Chloropidae *Hippelates* spp. | 0 | 1 | 0.01 | 1.0000 |
| Woodland | Chloropidae *Incertella* spp. | 1 | 0 | 0.33 | 0.0546 |
| Woodland | Dolichopodidae *Asyndetus* spp. | 0 | 1 | 0.17 | 0.3127 |
| Woodland | Dolichopodidae *Chrysotus* spp. | 1 | 0 | 0.02 | 1.0000 |
| Woodland | Dolichopodidae *Condylostylus* spp. | 1 | 0 | 0.65 | 0.0001 |
| Woodland | Fannidae *Fannia pusio* | 1 | 0 | 0.15 | 1.0000 |
| Woodland | Hybotidae Tachydromiinae spp. | 1 | 0 | 0.27 | 0.2297 |
| Woodland | Muscidae *Atherigona orientalis* | 1 | 0 | 0.05 | 1.0000 |

| Landscape measure | Taxon | Less bare ground | More bare ground | Test statistic | *P*-value |
|---|---|---|---|---|---|
| Woodland | Muscidae *Caricea erythrocera* | 1 | 0 | 0.22 | 0.4976 |
| Woodland | Muscidae *Polietes* sp. 1 | 1 | 0 | 0.09 | 1.0000 |
| Woodland | Muscidae spp. | 0 | 1 | 0.01 | 1.0000 |
| Woodland | Phoridae Group 1 | 1 | 0 | 0.21 | 0.3467 |
| Woodland | Phoridae Group 2 | 1 | 0 | 0.18 | 0.3313 |
| Woodland | Pipunculidae *Tomosvaryella* spp. | 0 | 1 | 0.01 | 1.0000 |
| Woodland | Sarcophagidae *Argoravinia modesta* | 1 | 0 | 0.21 | 0.3476 |
| Woodland | Sarcophagidae *Bercaea cruentata* | 1 | 0 | 0.22 | 0.4919 |
| Woodland | Sarcophagidae *Comasarcophaga* spp. | 1 | 0 | 0.09 | 1.0000 |
| Woodland | Sarcophagidae *Helicobia rapax* | 1 | 0 | 0.43 | 0.0103 |
| Woodland | Sarcophagidae Miltogramminae spp. | 1 | 0 | 0.32 | 0.1058 |
| Woodland | Sarcophagidae *Ravinia* spp. | 1 | 0 | 0.22 | 0.2361 |
| Woodland | Sarcophagidae *Tylomyia* spp. | 1 | 0 | 0.09 | 1.0000 |
| Woodland | Sciaridae *Eugnoriste brevirostris* | 1 | 0 | 0.15 | 1.0000 |
| Woodland | Sciaridae spp. | 1 | 0 | 0.36 | 0.0469 |
| Woodland | Stratiomyidae *Nemotelus kansensis* | 0 | 1 | 0.01 | 1.0000 |
| Woodland | Syrphidae *Copestylum tamaulipanum* | 0 | 1 | 0.01 | 1.0000 |
| Woodland | Syrphidae *Palpada vinetorum* | 1 | 0 | 0.22 | 0.4828 |
| Woodland | Syrphidae *Toxomerus marginatus* | 0 | 1 | 0.10 | 0.6011 |
| Woodland | Tachinidae *Cylindromyia intermedia* | 0 | 1 | 0.08 | 0.6636 |
| Woodland | Tachinidae *Phasia* spp. | 1 | 0 | 0.22 | 0.4874 |

**Table S20:** Test statistics for indicator species analyses investigating impacts of landscape-scale woodland cover on Diptera community composition. "1" values indicate which habitat category each taxon is associated with.

| **Taxon** | **Less woodland** | **More woodland** | **Test statistic** | ***P*-value** |
|---|---|---|---|---|
| Asilidae *Mallophora orcina* | 1 | 0 | 0.22 | 0.4819 |
| Asilidae *Ospriocerus* spp. | 0 | 1 | 0.10 | 0.6085 |
| Asilidae *Psilocurus* spp. | 1 | 0 | 0.22 | 0.4831 |
| Asteiidae *Leiomyza* spp. | 1 | 0 | 0.27 | 0.2327 |
| Bombyliidae *Chrysanthrax cypris* | 0 | 1 | 0.20 | 0.2454 |
| Bombyliidae *Chrysanthrax edititius* | 1 | 0 | 0.22 | 0.4777 |
| Bombyliidae *Geron* spp. | 0 | 1 | 0.44 | 0.0061 |
| Bombyliidae *Hemipenthes sinuosa* | 0 | 1 | 0.01 | 1.0000 |
| Bombyliidae *Poecilanthrax lucifer* | 0 | 1 | 0.22 | 0.2334 |
| Bombyliidae *Poecilognathus punctipennis* | 0 | 1 | 0.28 | 0.1128 |
| Bombyliidae *Poecilognathus* sp. 1 | 1 | 0 | 0.18 | 0.4146 |
| Bombyliidae *Rhynchanthrax* spp. | 0 | 1 | 0.22 | 0.2349 |
| Bombyliidae *Systoechus solitus* | 1 | 0 | 0.09 | 1.0000 |
| Bombyliidae *Villa lateralis* | 0 | 1 | 0.01 | 1.0000 |
| Calliphoridae *Lucilia* spp. | 0 | 1 | 0.22 | 0.2308 |
| Chloropidae *Apallates* spp. | 1 | 0 | 0.31 | 0.1014 |
| Chloropidae *Chlorops* spp. | 0 | 1 | 0.30 | 0.0691 |
| Chloropidae *Conioscinella* spp. | 0 | 1 | 0.22 | 0.1841 |
| Chloropidae *Hippelates* spp. | 1 | 0 | 0.22 | 0.4972 |
| Chloropidae *Incertella* spp. | 1 | 0 | 0.03 | 1.0000 |
| Dolichopodidae *Asyndetus* spp. | 1 | 0 | 0.25 | 0.1626 |
| Dolichopodidae *Chrysotus* spp. | 0 | 1 | 0.07 | 0.7729 |
| Dolichopodidae *Condylostylus* spp. | 1 | 0 | 0.26 | 0.1164 |

| **Taxon** | **Less woodland** | **More woodland** | **Test statistic** | ***P*-value** |
|---|---|---|---|---|
| Fannidae *Fannia pusio* | 1 | 0 | 0.15 | 1.0000 |
| Hybotidae Tachydromiinae spp. | 1 | 0 | 0.27 | 0.2326 |
| Muscidae *Atherigona orientalis* | 1 | 0 | 0.43 | 0.0103 |
| Muscidae *Caricea erythrocera* | 0 | 1 | 0.01 | 1.0000 |
| Muscidae *Polietes* sp. 1 | 1 | 0 | 0.09 | 1.0000 |
| Muscidae spp. | 0 | 1 | 0.01 | 1.0000 |
| Phoridae Group 1 | 1 | 0 | 0.06 | 1.0000 |
| Phoridae Group 2 | 1 | 0 | 0.18 | 0.3357 |
| Pipunculidae *Tomosvaryella* spp. | 1 | 0 | 0.32 | 0.1100 |
| Sarcophagidae *Argoravinia modesta* | 1 | 0 | 0.36 | 0.0476 |
| Sarcophagidae *Bercaea cruentata* | 1 | 0 | 0.22 | 0.4877 |
| Sarcophagidae *Comasarcophaga* spp. | 1 | 0 | 0.27 | 0.2310 |
| Sarcophagidae *Helicobia rapax* | 1 | 0 | 0.32 | 0.0706 |
| Sarcophagidae Miltogramminae spp. | 0 | 1 | 0.01 | 1.0000 |
| Sarcophagidae *Ravinia* spp. | 1 | 0 | 0.22 | 0.2289 |
| Sarcophagidae *Tylomyia* spp. | 1 | 0 | 0.27 | 0.2317 |
| Sciaridae *Eugnoriste brevirostris* | 1 | 0 | 0.15 | 1.0000 |
| Sciaridae spp. | 1 | 0 | 0.06 | 1.0000 |
| Stratiomyidae *Nemotelus kansensis* | 0 | 1 | 0.22 | 0.2346 |
| Syrphidae *Copestylum tamaulipanum* | 1 | 0 | 0.15 | 0.6004 |
| Syrphidae *Palpada vinetorum* | 0 | 1 | 0.01 | 1.0000 |
| Syrphidae *Toxomerus marginatus* | 1 | 0 | 0.27 | 0.2347 |
| Tachinidae *Cylindromyia intermedia* | 1 | 0 | 0.21 | 0.3421 |
| Tachinidae *Phasia* spp. | 0 | 1 | 0.22 | 0.2326 |

**Table S21:** Test statistics for indicator species analyses investigating impacts of **local floral richness and landscape-scale woodland cover** on Diptera and Hymenoptera community composition. "1" values indicate which habitat category each taxon is associated with.

| **Order** | **Taxon** | **Few flowers, little woodland** | **Few flowers, more woodland** | **More flowers, little woodland** | **More flowers, more woodland** | **Test statistic** | ***P*-value** |
|---|---|---|---|---|---|---|---|
| Diptera | Asilidae *Mallophora orcina* | 0 | 0 | 1 | 0 | 0.42 | 0.0800 |
| Diptera | Asilidae *Ospriocerus* spp. | 0 | 1 | 1 | 1 | 0.17 | 0.6980 |
| Diptera | Asilidae *Psilocurus* spp. | 1 | 0 | 0 | 0 | 0.35 | 0.2355 |
| Diptera | Asteiidae *Leiomyza* spp. | 1 | 0 | 0 | 0 | 0.43 | 0.0529 |
| Diptera | Bombyliidae *Chrysanthrax cypris* | 1 | 1 | 0 | 1 | 0.25 | 0.4986 |
| Diptera | Bombyliidae *Chrysanthrax edititius* | 1 | 0 | 1 | 0 | 0.22 | 0.8238 |
| Diptera | Bombyliidae *Geron* spp. | 0 | 1 | 0 | 1 | 0.44 | 0.0252 |
| Diptera | Bombyliidae *Hemipenthes sinuosa* | 1 | 0 | 1 | 1 | 0.18 | 0.7622 |
| Diptera | Bombyliidae *Poecilanthrax lucifer* | 0 | 0 | 0 | 1 | 0.36 | 0.1557 |
| Diptera | Bombyliidae *Poecilognathus punctipennis* | 0 | 1 | 0 | 1 | 0.27 | 0.3489 |
| Diptera | Bombyliidae *Poecilognathus* sp. 1 | 1 | 0 | 0 | 0 | 0.26 | 0.3492 |
| Diptera | Bombyliidae *Rhynchanthrax* spp. | 0 | 1 | 0 | 0 | 0.42 | 0.0814 |
| Diptera | Bombyliidae *Systoechus solitus* | 1 | 0 | 1 | 1 | 0.16 | 1.0000 |
| Diptera | Bombyliidae *Villa lateralis* | 1 | 0 | 1 | 1 | 0.18 | 0.7588 |
| Diptera | Calliphoridae *Lucilia* spp. | 0 | 0 | 0 | 1 | 0.36 | 0.1464 |
| Diptera | Chloropidae *Apallates* spp. | 1 | 0 | 1 | 0 | 0.31 | 0.1861 |
| Diptera | Chloropidae *Chlorops* spp. | 0 | 1 | 0 | 1 | 0.32 | 0.1499 |

| Order | Taxon | Few flowers, little woodland | Few flowers, more woodland | More flowers, little woodland | More flowers, more woodland | Test statistic | *P*-value |
|---|---|---|---|---|---|---|---|
| Diptera | Chloropidae *Conioscinella* spp. | 0 | 1 | 0 | 1 | 0.21 | 0.5872 |
| Diptera | Chloropidae *Hippelates* spp. | 1 | 0 | 0 | 0 | 0.35 | 0.2407 |
| Diptera | Chloropidae *Incertella* spp. | 0 | 0 | 1 | 0 | 0.11 | 0.9430 |
| Diptera | Dolichopodidae *Asyndetus* spp. | 0 | 0 | 1 | 0 | 0.51 | 0.0071 |
| Diptera | Dolichopodidae *Chrysotus* spp. | 1 | 1 | 0 | 1 | 0.40 | 0.0681 |
| Diptera | Dolichopodidae *Condylostylus* spp. | 1 | 0 | 1 | 1 | 0.35 | 0.1452 |
| Diptera | Fannidae *Fannia pusio* | 0 | 0 | 1 | 0 | 0.29 | 0.4269 |
| Diptera | Hybotidae Tachydromiinae spp. | 0 | 0 | 1 | 0 | 0.52 | 0.0131 |
| Diptera | Muscidae *Atherigona orientalis* | 1 | 0 | 1 | 0 | 0.42 | 0.0296 |
| Diptera | Muscidae *Caricea erythrocera* | 0 | 0 | 1 | 1 | 0.23 | 0.5721 |
| Diptera | Muscidae *Polietes* sp. 1 | 1 | 0 | 0 | 1 | 0.25 | 0.6086 |
| Diptera | Muscidae spp. | 1 | 1 | 0 | 0 | 0.22 | 0.8280 |
| Diptera | Phoridae Group 1 | 1 | 0 | 0 | 0 | 0.23 | 0.4551 |
| Diptera | Phoridae Group 2 | 1 | 0 | 1 | 1 | 0.25 | 0.4685 |
| Diptera | Pipunculidae *Tomosvaryella* spp. | 0 | 0 | 1 | 0 | 0.44 | 0.0311 |
| Diptera | Sarcophagidae *Argoravinia modesta* | 0 | 0 | 1 | 0 | 0.37 | 0.0776 |
| Diptera | Sarcophagidae *Bercaea cruentata* | 1 | 0 | 1 | 0 | 0.22 | 0.8307 |
| Diptera | Sarcophagidae *Comasarcophaga* spp. | 1 | 0 | 1 | 0 | 0.27 | 0.4686 |
| Diptera | Sarcophagidae *Helicobia rapax* | 1 | 0 | 1 | 0 | 0.33 | 0.1545 |
| Diptera | Sarcophagidae Miltogramminae spp. | 1 | 0 | 0 | 1 | 0.30 | 0.3149 |
| Diptera | Sarcophagidae *Ravinia* spp. | 1 | 0 | 1 | 0 | 0.23 | 0.5691 |

| Order | Taxon | Few flowers, little woodland | Few flowers, more woodland | More flowers, little woodland | More flowers, more woodland | Test statistic | *P*-value |
|---|---|---|---|---|---|---|---|
| Diptera | Sarcophagidae *Tylomyia* spp. | 1 | 0 | 1 | 0 | 0.27 | 0.4642 |
| Diptera | Sciaridae *Eugnoriste brevirostris* | 0 | 0 | 1 | 0 | 0.29 | 0.4125 |
| Diptera | Sciaridae spp. | 0 | 0 | 1 | 0 | 0.34 | 0.1860 |
| Diptera | Stratiomyidae *Nemotelus kansensis* | 0 | 1 | 0 | 1 | 0.23 | 0.5648 |
| Diptera | Syrphidae *Copestylum tamaulipanum* | 0 | 0 | 1 | 0 | 0.25 | 0.4187 |
| Diptera | Syrphidae *Palpada vinetorum* | 1 | 0 | 0 | 1 | 0.20 | 1.0000 |
| Diptera | Syrphidae *Toxomerus marginatus* | 1 | 0 | 1 | 0 | 0.27 | 0.4599 |
| Diptera | Tachinidae *Cylindromyia intermedia* | 1 | 1 | 1 | 0 | 0.22 | 0.6287 |
| Diptera | Tachinidae *Phasia* spp. | 0 | 0 | 0 | 1 | 0.36 | 0.1628 |
| Hymenoptera | Andrenidae *Andrena rudbeckiae* | 1 | 1 | 0 | 1 | 0.16 | 1.0000 |
| Hymenoptera | Andrenidae *Calliopsis andreniformis* | 1 | 0 | 1 | 0 | 0.32 | 0.2020 |
| Hymenoptera | Andrenidae *Perdita cambarella* | 0 | 0 | 1 | 0 | 0.32 | 0.1324 |
| Hymenoptera | Andrenidae *Perdita ignota* | 1 | 0 | 0 | 1 | 0.20 | 1.0000 |
| Hymenoptera | Apidae *Apis mellifera* | 0 | 0 | 1 | 1 | 0.40 | 0.0653 |
| Hymenoptera | Apidae *Bombus fraternus* | 0 | 0 | 0 | 1 | 0.45 | 0.0301 |
| Hymenoptera | Apidae *Bombus griseocollis* | 1 | 0 | 1 | 1 | 0.19 | 0.6753 |
| Hymenoptera | Apidae *Bombus pensylvanicus* | 1 | 0 | 0 | 1 | 0.25 | 0.4149 |
| Hymenoptera | Apidae *Centris atripes* | 0 | 0 | 1 | 1 | 0.23 | 0.5691 |
| Hymenoptera | Apidae *Ceratina shinnersi* | 1 | 0 | 0 | 0 | 0.24 | 1.0000 |
| Hymenoptera | Apidae *Ceratina strenua* | 1 | 0 | 1 | 1 | 0.18 | 0.8995 |
| Hymenoptera | Apidae *Diadasia enavata* | 0 | 1 | 0 | 1 | 0.23 | 0.5778 |

| Order | Taxon | Few flowers, little woodland | Few flowers, more woodland | More flowers, little woodland | More flowers, more woodland | Test statistic | *P*-value |
|---|---|---|---|---|---|---|---|
| Hymenoptera | Apidae *Diadasia rinconis* | 0 | 1 | 0 | 1 | 0.23 | 0.5685 |
| Hymenoptera | Apidae *Melissodes communis* | 1 | 1 | 1 | 0 | 0.40 | 0.0487 |
| Hymenoptera | Apidae *Melissodes coreopsis* | 0 | 1 | 0 | 0 | 0.17 | 0.8492 |
| Hymenoptera | Apidae *Melissodes tepaneca* | 0 | 0 | 1 | 0 | 0.55 | 0.0026 |
| Hymenoptera | Apidae *Melissodes wheeleri* | 1 | 0 | 1 | 0 | 0.22 | 0.8313 |
| Hymenoptera | Apidae *Epimelissodes obliqua* | 0 | 1 | 1 | 0 | 0.24 | 0.3285 |
| Hymenoptera | Apidae *Epimelissodes petulca* | 0 | 1 | 0 | 0 | 0.44 | 0.0312 |
| Hymenoptera | Apidae *Xylocopa virginica* | 0 | 1 | 1 | 1 | 0.45 | 0.0159 |
| Hymenoptera | Braconidae sp. 2 | 0 | 0 | 0 | 1 | 0.26 | 0.5273 |
| Hymenoptera | Colletidae *Colletes mandibularis* | 0 | 0 | 1 | 1 | 0.23 | 0.5639 |
| Hymenoptera | Crabronidae *Astata clypeata* | 0 | 0 | 1 | 1 | 0.23 | 0.5748 |
| Hymenoptera | Crabronidae *Liris argentatus* | 0 | 0 | 1 | 0 | 0.32 | 0.2006 |
| Hymenoptera | Crabronidae *Soleriella* sp. 1 | 0 | 1 | 1 | 0 | 0.26 | 0.3882 |
| Hymenoptera | Crabronidae *Soleriella* sp. 2 | 0 | 1 | 0 | 0 | 0.29 | 0.4148 |
| Hymenoptera | Crabronidae *Tachysphex* sp. 1 | 0 | 1 | 1 | 0 | 0.30 | 0.2537 |
| Hymenoptera | Crabronidae *Tachysphex* sp. 2 | 0 | 1 | 1 | 1 | 0.15 | 0.8089 |
| Hymenoptera | Crabronidae *Tachysphex* sp. 3 | 1 | 1 | 0 | 0 | 0.22 | 0.8269 |
| Hymenoptera | Crabronidae *Tachysphex* sp. 6 | 0 | 1 | 0 | 1 | 0.24 | 0.3753 |
| Hymenoptera | Crabronidae *Tachytes amazonus* | 0 | 1 | 0 | 1 | 0.23 | 0.5707 |
| Hymenoptera | Crabronidae *Trypoxylon* sp. 1 | 1 | 1 | 1 | 0 | 0.19 | 0.6353 |
| Hymenoptera | Halictidae *Agapostemon splendens* | 0 | 0 | 1 | 1 | 0.23 | 0.5702 |
| Hymenoptera | Halictidae *Agapostemon texanus* | 0 | 0 | 1 | 1 | 0.29 | 0.2237 |
| Hymenoptera | Halictidae *Augochlorella aurata* | 1 | 0 | 0 | 0 | 0.53 | 0.0032 |

| Order | Taxon | Few flowers, little woodland | Few flowers, more woodland | More flowers, little woodland | More flowers, more woodland | Test statistic | *P*-value |
|---|---|---|---|---|---|---|---|
| Hymenoptera | Halictidae *Augochloropsis metallica* | 1 | 0 | 1 | 1 | 0.22 | 0.6270 |
| Hymenoptera | Halictidae *Augochloropsis sumptuosa* | 1 | 0 | 0 | 1 | 0.20 | 1.0000 |
| Hymenoptera | Halictidae *Halictus ligatus* | 0 | 0 | 1 | 0 | 0.34 | 0.1777 |
| Hymenoptera | Halictidae *Lasioglossum bardum* | 0 | 1 | 0 | 0 | 0.29 | 0.4148 |
| Hymenoptera | Halictidae *Lasioglossum bruneri* | 0 | 0 | 1 | 1 | 0.19 | 0.6693 |
| Hymenoptera | Halictidae *Lasioglossum callidum* | 0 | 1 | 0 | 1 | 0.23 | 0.5730 |
| Hymenoptera | Halictidae *Lasioglossum coactus* | 0 | 0 | 1 | 1 | 0.17 | 0.7935 |
| Hymenoptera | Halictidae *Lasioglossum connexum* | 0 | 1 | 0 | 1 | 0.25 | 0.4127 |
| Hymenoptera | Halictidae *Lasioglossum coreopsis* | 1 | 0 | 0 | 0 | 0.41 | 0.0377 |
| Hymenoptera | Halictidae *Lasioglossum disparile* | 0 | 0 | 1 | 1 | 0.21 | 0.5357 |
| Hymenoptera | Halictidae *Lasioglossum fedorense* | 0 | 0 | 1 | 0 | 0.29 | 0.4134 |
| Hymenoptera | Halictidae *Lasioglossum hitchensi* | 0 | 0 | 1 | 0 | 0.17 | 0.8419 |
| Hymenoptera | Halictidae *Lasioglossum hudsoniellum* | 1 | 1 | 1 | 0 | 0.16 | 0.7707 |
| Hymenoptera | Halictidae *Lasioglossum hunteri* | 0 | 0 | 1 | 1 | 0.23 | 0.5616 |
| Hymenoptera | Halictidae *Lasioglossum illinoense* | 0 | 0 | 0 | 1 | 0.32 | 0.2420 |
| Hymenoptera | Halictidae *Lasioglossum lustrans* | 1 | 0 | 1 | 0 | 0.22 | 0.8272 |
| Hymenoptera | Halictidae *Lasioglossum pectorale* | 1 | 0 | 1 | 0 | 0.22 | 0.8272 |

| Order | Taxon | Few flowers, little woodland | Few flowers, more woodland | More flowers, little woodland | More flowers, more woodland | Test statistic | *P*-value |
|---|---|---|---|---|---|---|---|
| Hymenoptera | Halictidae *Lasioglossum* sp. OK-1 | 1 | 0 | 1 | 0 | 0.22 | 0.8247 |
| Hymenoptera | Halictidae *Lasioglossum* sp. TX-24 | 0 | 1 | 0 | 0 | 0.09 | 1.0000 |
| Hymenoptera | Halictidae *Lasioglossum* sp. TX-3 | 0 | 1 | 1 | 0 | 0.32 | 0.1869 |
| Hymenoptera | Halictidae *Lasioglossum* sp. TX-6 | 0 | 0 | 0 | 1 | 0.26 | 0.5310 |
| Hymenoptera | Halictidae *Lasioglossum tegulare* | 1 | 0 | 1 | 0 | 0.28 | 0.3238 |
| Hymenoptera | Megachilidae *Anthidiellum notatum* | 0 | 1 | 0 | 1 | 0.23 | 0.5719 |
| Hymenoptera | Megachilidae *Dianthidium curvatum* | 0 | 0 | 0 | 1 | 0.32 | 0.2417 |
| Hymenoptera | Megachilidae *Dianthidium texanum* | 0 | 1 | 1 | 0 | 0.30 | 0.2583 |
| Hymenoptera | Megachilidae *Heriades carinata* | 0 | 1 | 0 | 0 | 0.29 | 0.4190 |
| Hymenoptera | Megachilidae *Megachile albitarsis* | 1 | 1 | 0 | 1 | 0.18 | 0.7583 |
| Hymenoptera | Megachilidae *Megachile brevis* | 1 | 0 | 0 | 1 | 0.22 | 0.4537 |
| Hymenoptera | Megachilidae *Megachile comata* | 0 | 1 | 0 | 1 | 0.27 | 0.3434 |
| Hymenoptera | Megachilidae *Megachile inimica* | 0 | 0 | 0 | 1 | 0.25 | 0.6959 |
| Hymenoptera | Megachilidae *Megachile mendica* | 1 | 1 | 0 | 1 | 0.16 | 1.0000 |
| Hymenoptera | Megachilidae *Megachile montivaga* | 1 | 0 | 0 | 0 | 0.23 | 0.4570 |
| Hymenoptera | Megachilidae *Megachile parallela* | 0 | 1 | 1 | 1 | 0.26 | 0.3434 |

| Order | Taxon | Few flowers, little woodland | Few flowers, more woodland | More flowers, little woodland | More flowers, more woodland | Test statistic | *P*-value |
|---|---|---|---|---|---|---|---|
| Hymenoptera | Megachilidae *Megachile policaris* | 0 | 1 | 1 | 0 | 0.32 | 0.1373 |
| Hymenoptera | Megachilidae *Osmia chalybea* | 0 | 1 | 0 | 1 | 0.23 | 0.5696 |
| Hymenoptera | Megachilidae *Osmia subfasciata* | 0 | 0 | 1 | 0 | 0.44 | 0.0322 |
| Hymenoptera | Pompilidae *Ageniella accepta* | 0 | 1 | 0 | 1 | 0.27 | 0.3492 |
| Hymenoptera | Pompilidae *Ageniella arcuata* | 0 | 0 | 0 | 1 | 0.26 | 0.3030 |
| Hymenoptera | Pompilidae *Ageniella placita* | 0 | 1 | 1 | 0 | 0.35 | 0.0902 |
| Hymenoptera | Pompilidae *Anoplius nigritus* | 0 | 0 | 1 | 0 | 0.52 | 0.0135 |
| Hymenoptera | Pompilidae *Anoplius* sp. 3 | 0 | 0 | 1 | 0 | 0.44 | 0.0299 |
| Hymenoptera | Pompilidae *Aporinellus yucatanensis* | 0 | 0 | 1 | 1 | 0.37 | 0.1255 |
| Hymenoptera | Scelionidae sp. g | 0 | 0 | 1 | 0 | 0.29 | 0.4176 |
| Hymenoptera | Sphecidae *Sceliphron caementarium* | 1 | 0 | 0 | 0 | 0.24 | 1.0000 |
| Hymenoptera | Tiphiidae *Myzinum maculata* | 0 | 0 | 0 | 1 | 0.32 | 0.2348 |
| Hymenoptera | Tiphiidae *Myzinum quinquecinctum* | 1 | 0 | 1 | 1 | 0.16 | 1.0000 |
| Hymenoptera | Vespidae *Parancistrocerus fulvipes* | 0 | 0 | 0 | 1 | 0.25 | 0.6981 |
| Hymenoptera | Vespidae *Polistes bellicosus* | 0 | 0 | 1 | 0 | 0.30 | 0.2466 |
| Hymenoptera | Vespidae *Polistes carolina* | 1 | 0 | 0 | 1 | 0.20 | 1.0000 |
| Hymenoptera | Vespidae *Polistes dorsalis* | 0 | 1 | 1 | 0 | 0.33 | 0.1181 |
| Hymenoptera | Vespidae *Polistes fuscatus* | 1 | 0 | 1 | 0 | 0.22 | 0.8248 |
| Hymenoptera | Vespidae *Stenodynerus propinquus* | 0 | 0 | 1 | 1 | 0.27 | 0.3453 |

**Figure S1: Impacts of (a) bare ground, (b) landscape-scale woodland cover, and (c,d) local floral richness and woodland cover on Diptera (a-c) and Hymenoptera (d) community composition. Plots show principal coordinate analysis ordinations, with group centroids and standard deviation data ellipses.**

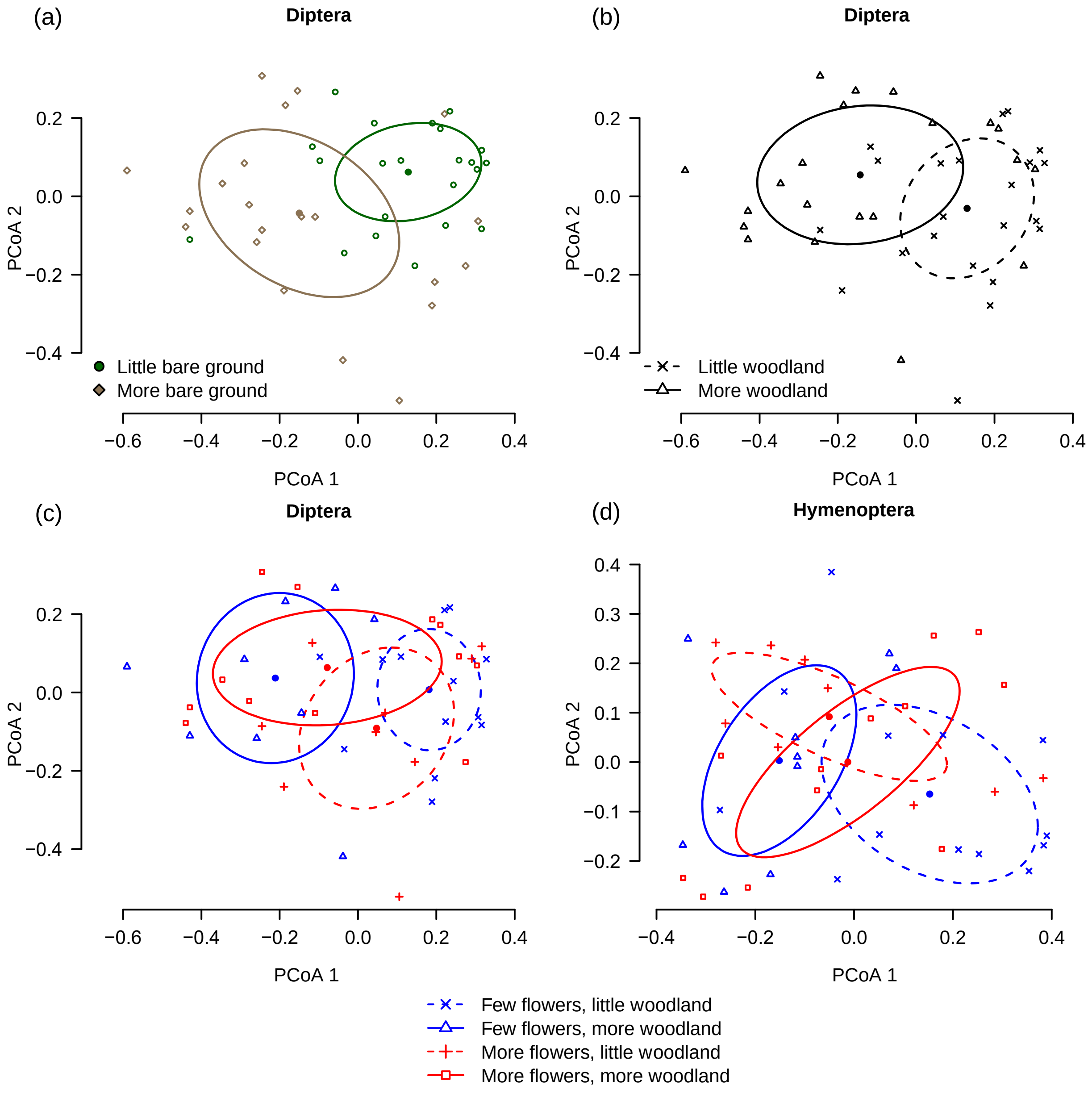
